# Bottom-up control of leakage in spectral electrophysiological source imaging via structured sparse bayesian learning

**DOI:** 10.1101/2020.02.25.964684

**Authors:** Eduardo Gonzalez-Moreira, Deirel Paz-Linares, Ariosky Areces-Gonzalez, Ying Wang, Min Li, Thalia Harmony, Pedro A. Valdes-Sosa

## Abstract

Brain electrical activity in different spectral bands has been associated with diverse mechanisms underlying Brain function. Deeper reconnoitering of these mechanisms entails mapping in grayordinates (Gray Matter coordinates), the spectral features of electrophysiological Brain signals. Such mapping is possible through MEG/EEG signals, due to their wide Brain coverage and excellent temporal resolution in reflecting neural-electrical-activity. This process-coined Electrophysiological Source Imaging (ESI)-can only produce approximated images of Brain activity, which are severely distorted by leakage: a pervasive effect in almost any imaging technique. It has been proposed that leakage control to tolerable levels can be achived through using priors or regularization within ESI, but their implementation commonly yields meager statistical guaranties. We introduce bottom-up control of leakage: defined as maximum Bayesian evidence search braced with priors precisely on the spectral responses. This is feasible due to an instance of Bayesian learning of complex valued data: spectral Structured Sparse Bayesian Learning (sSSBL). “Spectral” refers to specific spatial topologies that are reflected by the MEG/EEG spectra. We also present a new validation benchmark based on the concurrency between high density MEG and its associated pseudo-EEG of lower density. This reveals that prevealing methods like eLORETA and LCMV can fall short of expectations whereas sSSBL exibits an exellent performance. A final qualitative assesment reveals that sSSBL can outline brain lessions using just low density EEG, according to the T2 MRI shine through of the affected areas.

## Introduction

Band specific synchronized-spectral-activity is the underlying mechanism for large scale Brain integration or networks of functionally specialized regions from which coherent behavior and cognition emerge [Engel et al., 2001; Varela et al. 2001; Valdes-Sosa et al 1992; Nunez and Srinivasan 2006; Vidaurre et al 2018]. Its cause dwells mostly microscopically in the neural architecture and interactions within layers at the cortical columns [Freeman 1975; Jirsa 1997], but still macroscopically its topological organization-spatial distribution and connectivity pattern-at the macroscopic level possesses a tremendous descriptive power in behavioral and clinical aspects of healthy, developing, aging, and diseased human brains [Bullmore and Sporns 2009; Rubinov and Sporns 2010; Sporns 2011].

An important part in describing this spectral activity involves mapping its spatial distribution alone, by the localization of responsive areas at the observational level of experimental techniques like Functional Magnetic Resonance Imaging (fMRI) and Electro/Magnetoencephalogram (EEG/MEG) [Mantini 2007]. For either technique, mapping these responses cannot be tackled directly from the observed data: the spectral composition of fMRI signals is severely distorted (spectral leakage) by the slow metabolic-hemodynamic cascade of processes following the actual neural activations [Buxton 1998; Logothetis 2001], whereas EEG/MEG signals are affected in a different manner, their low resolution and blurring effect of head volume conduction (activation leakage) [Haufe et al 2013, Stokes et al 2017, Van de Steen et al 2019].

However, MEG/EEG signals have purely electrophysiological causes. Synchronic neural activity, at both microscopic and macroscopic Brain scales, elicit observable currents that are sensed through EEG electrodes or MEG gradiometers. Therefore, these signals are a sharper reflection of neural activity ***ι***(*t*), converted directly into voltage ***v***(*t*) or magnetic field ***b***(*t*) by the laws of the electromagnetic field in media alone. In the neuroimaging context this is denominated forward equation, a linear model whose operator is coined with the term Lead Field (LF) **L**_***vι***_ [Grech et al 2008; Nunez et al 1994, Burle et al 2015], see Figure1. Full details on the notation in Section 1 of the “Supporting Information” (SI) SI-1. Thus, EEG/MEG carry on rich information of the neural mechanisms underpinned by purely electrical activity, that is registered with excellent temporal resolution and without spectral distortion. This is the reason why they have-initially EEG and later MEG-provided most of the existent knowledge on the healthy brain dynamics [Freeman 1975, Lopes da Silva 2013, Larson-Prior 2013].

**Figure 1:**
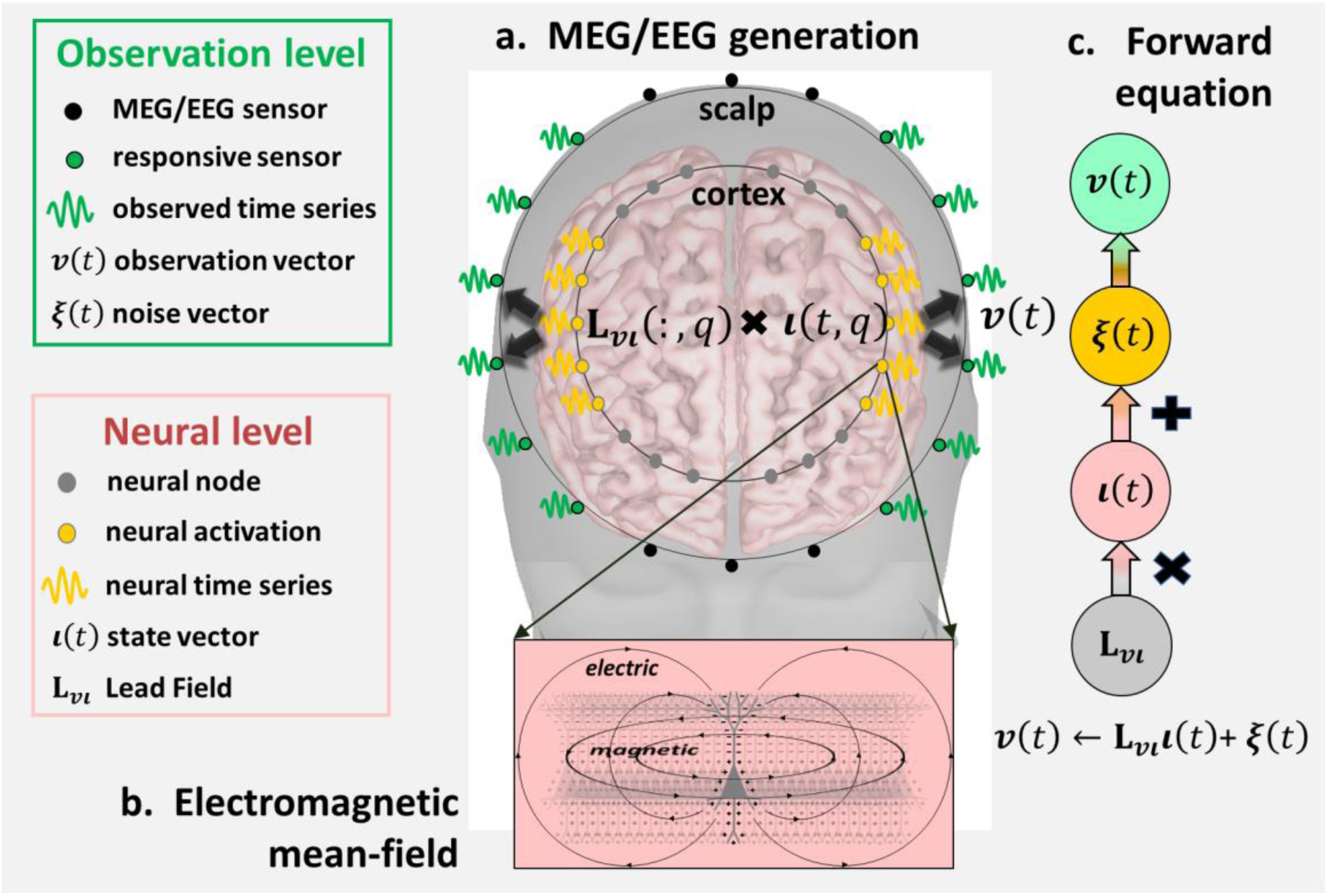
Illustration of the mechanisms underlying the generation of MEG/EEG signals. Their actual causes sojourn in the time variable (t) pattern of macroscopic currents at gray mater points (q) represented as **ι**(t, q). (a) these currents, originated from a homogeneous polarization of the pyramidal layers, sustain a local electromagnetic mean-field, and the spatial distribution of the external signal attributed to a single gray mater generator is explained in terms of a stationary linear equation for the electromagnetic field in media **L**_**vι**_(:, q). (b) **v**(t) can be observed externally through MEG/EEG sensors (electrodes or gradiometers). (c) from the principle of superposition stems the linear generative model of MEG/EEG signals explaining the composited effect of currents distributed across the whole gray mater **v**(t) ← **L**_**vι**_**ι**(t)+ **ξ**(t) where the process **ξ**(t) represents the effect of instrumental or environmental noise.

### Leakage of electrophysiological source imaging

The basic flaw is the pervasive activation leakage of routine Electrophysiological Source Imaging (ESI) solutions, pledged with the coarse spatial undertone of sensed EEG/MEG responses with Gray Matter (in grayordinates or Gray Matter coordinate system) activations [Hämäläinen et al 1994; Pascual-Marqui et al 1994; Pascual-Marqui 1999; Srinivasan 1999, Babiloni et al 2001, Pascual-Marqui 2002, He et al 2019]. Where ESI is defined as the pseudo-inversion of the operator **L**_***vι***_ of the Forward equation. This inversion process or ESI is formally expressed through a transfer operator **T**_***ιv***_ that attempts to explain in grayordinates the neural causes of signals [Nunez et al 1994, Burle et al 2015], see Figure 2.

**Figure 2:**
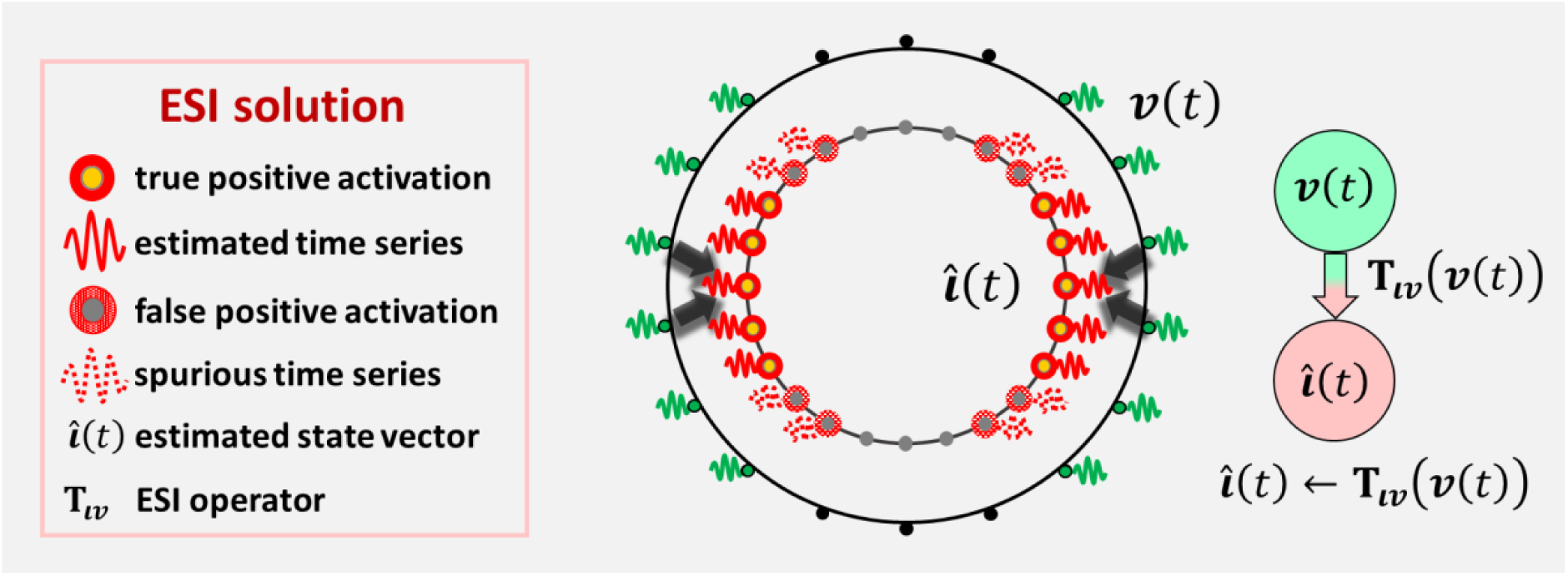
Basic leakage problem with ESI estimation. Attempting to retrieve the source activity via certain operator **T**_**ιv**_ (this can be linear/nonlinear and encode different prior information) typical methods produce a blurred estimation 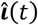. This carries on spurious (false positive) activations that extend beyond the truly active areas.

The pitfall of leakage has been followed by the deployment of several “generations” of ESI methods within just two decades [Gonzalez-Moreira et al 2018b]. The term “generation” coins basic differences of the statistical learning method towards the construction of the EEG/MEG source transfer operator -via Linear, Nonlinear Univariate, Nonlinear Multivariate models- and the inclusion of prior information in order to ameliorate Leakage, we defer details on this to SI-2.

More sophisticated models (Nonlinear Multivariate) leveraged by the 3^rd^ ESI generation (SI-2) can lead to very inaccurate results without adequate prior variable selection of 2^nd^ generation methods, that could possibly constrain the grayordinate active subspace. This flaw is caused by inaccurate high dimensional computations and ill-conditioning of the coactivation matrix (which regard multivariate patterns) used in their nonlinear algorithms [Friston et al 2007; Paz-Linares et al 2018; Gonzalez-Moreira et al 2018a; 2018b; 2018c]. Likewise, this is critical for EEG/MEG connectivity methods preceded by any ESI solution [Coulclogh, 2016; Marinazzo et al 2019; Haufe et al 2013, Stokes et al 2017, Van de Steen et al 2019]. Learning the truly responsive sources in the spectral domain remains an uneasy enterprise underscoring any further source space analysis. This is connected first, to the selection of priors for controlling leakage, second, statistical guarantees of the inference framework, and third, appropriate experimental confirmation.

### Bottom-up control of leakage

We aknowledge a major dificulty with leakage control in ESI arises from a very general methodological unconsitency, this is the top-down approach to a composited problem (Figure 3) in two steps, first a procedure targeting the solution to an ill posed problem at the top a) but which affects ireversibly derived statistical quantities intended to gauge a given effect at the bottom b). We are concerned with the case in which a) is stated as the discovery of hidden source activity from MEG/EEG observations in time or spectral domain via ESI (Figure 3a); and b) as the statistical relevance of spectral responses upon which the Leakage effect is ultimately observed. The latter are irrversibly affected by ESI inacurracy and therefore are a common target of Leakage corrections (Figure 3b).

**Figure 3:**
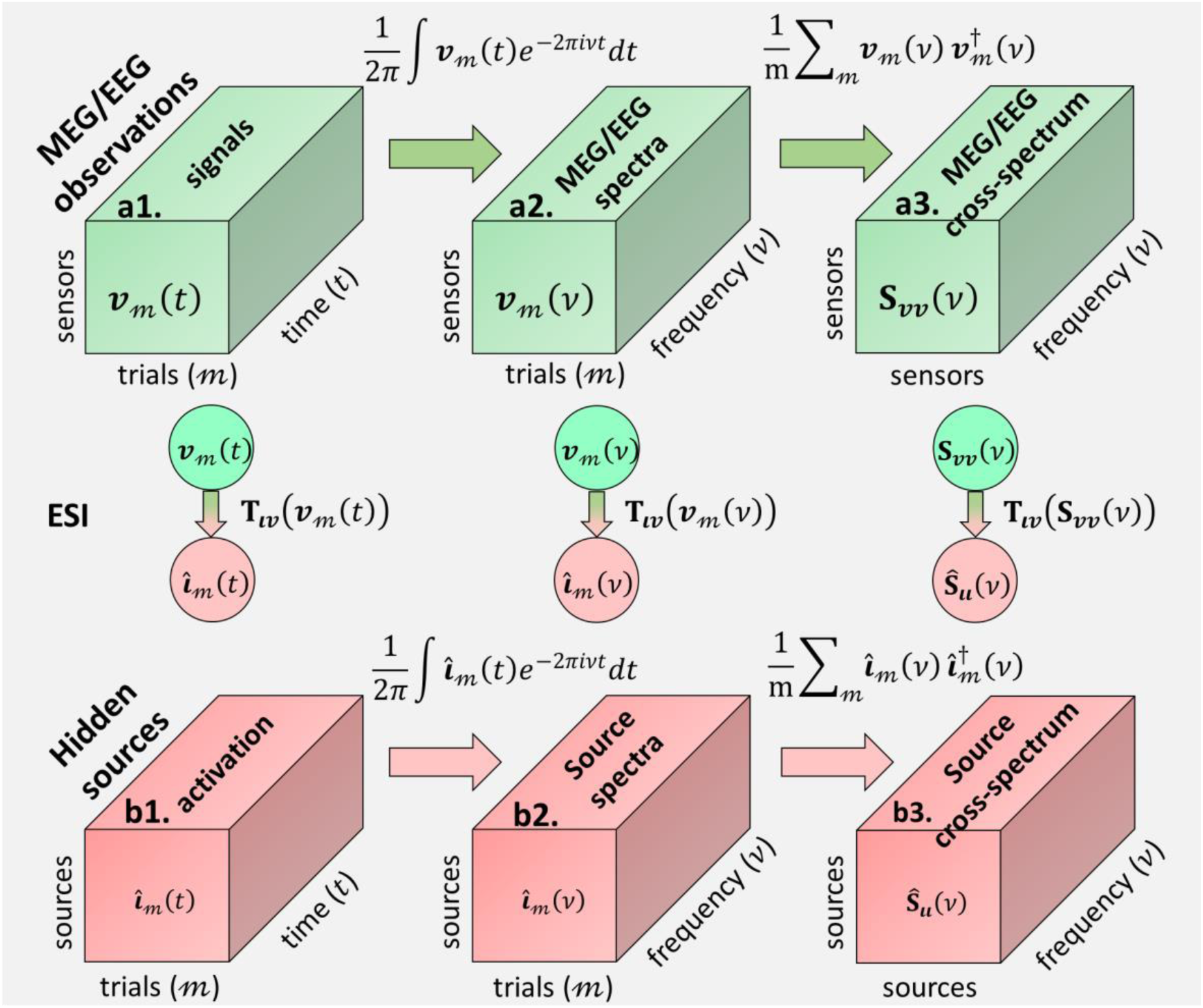
Transit from observations (a) to hidden source activation (b) via three possible ESI routes. The starting point is a tensor structure containing MEG/EEG signals (a1) that are recorded a long time (t) and for different trials (𝓂). These are used to discover spectral properties encoded by the cross-spectrum tensor (b3), i.e. the activations (tensor diagonal entries) and coactivation (tensor off-diagonal entries). From left to right, the ESI routes to define the source transfer operator stride across qualitatively different learning paths: via independent instances in time a1 → b1 and frequency a2 → b2 domains via the cross-spectrum a3 → b3.

As Figure 3 illustrates, the alternatives for solving leakage stride across three different ESI routes a_k_ → b_k_, k = 1,2,3, in the transit from observations a) to hidden source activation b). Note that the segregation of ESI routes refres to the form in which the data is used for statistical learning of the transfer operator **T**_***ιv***_ and in disregar to the type of solution which can be either Linear, Nonlinear univariated or Nonlinear multivariated. We enphasize that the top-down inconsistency relies on the lack of statistical guaranties in leakage control upon b_3_ via ESI routes a_1_ → b_1_ and a_2_ → b_2_, in other words, by means of priors on independent instances of source activity. Therefore, the only available alternative is bottom-up Leakage control by placing priors directly on the cross-spectrum b_3_ whose estimation is carried out via ESI route a_3_ → b_3_.

Indeed this explains why some implementions of 2^nd^ and 3^rd^ generation methods are far superior to the linear methods in the first generation. The reason relies on the idea that, within the nonlinear learning process, parameters underlaying the estimation across samples are prone to perform leakage control. An importamt limitation hitched to direct spectral ESI solutions is the unavailability of learning models for complex valued data. Aknowledging these facts, we present a spectral modality of ESI provided with the property of botom-up control of leakage. The target of our method is to directly estimate the cross-spectrum tensor b_3_, with complex valued structured sparsity priors via Bayesian learning. We refer to this method as Spectral Structured Sparse Bayesian Learning (sSSBL).

### Spectral Structured Sparse Bayesian Learning (sSSBL) and extensions (sSSBL++)

The basic principle stems behind the idea of EEG/MEG scalp-spectral features are caused by focalized cortical topologies and are therefore a better target for sparse ESI inversion (Figure 4a). sSSBL targets individual topologies by means of a group penalization model upon the sample space of spectral source reponses (***ι***_1_(*v*) … ***ι***_*𝓂*_(*v*)) that may possibly explain the observed cros-sspectum (**S**_***vv***_(*v*)) (Figure 4b). This translates into a sparse univariate model for the selection of rows/colums corresponding to diagonal elements on the source cross-spectrum (**S**_***ιι***_(*v*)), whose statistical relevance is controled locally through the variances 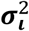 of the Hierarchical Elastic Net model. The global level of sparsity is controlled via a flexible twofolded regularization model with parameters (α, *ρ*) (Figure 4c). See the details on the formulation of the model underlying the sSSBL model in SI-3 and SI-4.

**Figure 4:**
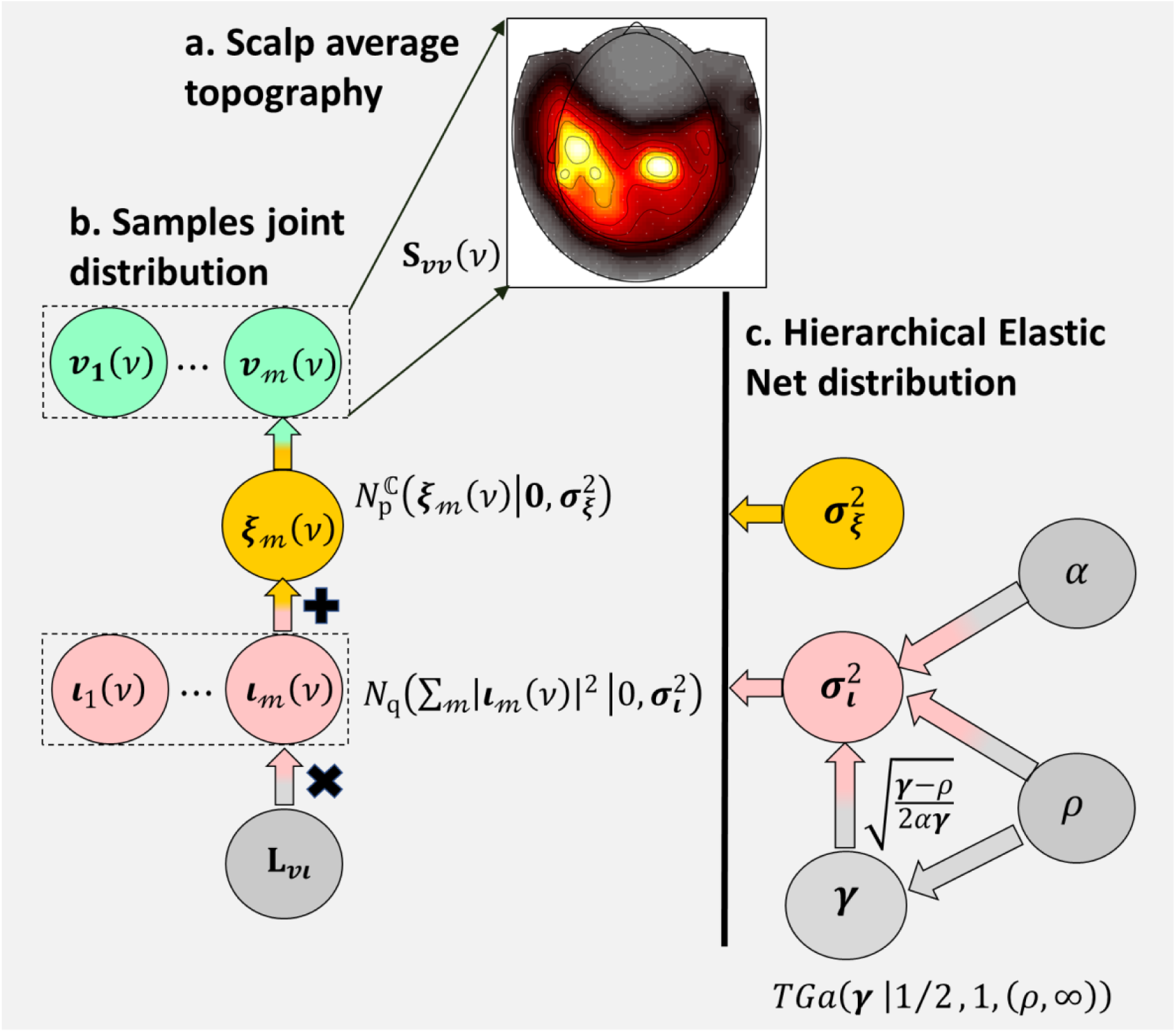
Model of the Spectral Structured Sparse Bayesian Learning. Priors are used directly on the spectral domain of the MEG/EEG generative model, to target source activity that explains a statistical tendency (a) at a given frequency of the observed cross-spectrum **S**_**vv**_(v). Gaussianity is assumed at two levels (b): First, on the spectral noise process **ξ**_𝓂_(v), as many typical ESI methods do in the time domain, which yields in this case a hermitian Gaussian model of the MEG/EEG cross-spectrum with variance spectral noise 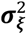. Second, on the spectral activations **ι**_𝓂_(v), but through a joint model of the different instances 𝓂. The statistical relevance of a given activation pattern common for all instances 𝓂 is determined by the variances 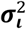. The estimation of these variances is based on the Hierarchical Elastic Net (c), with sparse variable selection performed through the parameters **γ** of Truncated Gamma distribution. The global level of sparsity or shrinkage is controlled by α and ρ.

The statistical gurantees in part rely on the maximum evidence towards determination of the variances that control leakage. Within the inference process these variances directly affect the source transfer operator **T**_***ιv***_, whose estimation is carried out here suboptimally, but highly eficientlly, via the Expectation-Maximization (E-M) algorithm. See the graphical sSSBL algorithm in the section 1 of “Materials and Methods” (MM) MM-1. With “suboptimal” we highlight the E-M limited capacity to only provide a local optima of the model evidence, but which can be higly accurate with the further inclusion of priors. We denominate this expanded sSSBL (sSSBL++). A critical set of five priors is introduced onto the model of Figure 4 serving to the purpose of not only constraining the optimal evidence search process but providing the estimated source activation with relevant physical features.

We use a joint activation field model, which introduces two different type of links (Figure 5a). First, a link onto its three spatial directions, via probabilistic 2D rotational invariance or isotropy (Figure 5b) reducing the degree of freedom, but still regarding surface normals as the field preferential direction. This diverges from the use of rigid constrains in assuming that surface normals are the only plausible orientation for the field of cortical currents. Second, a link onto the fields of neighbour cortical points via the Laplacian, which builds on assumptions of a graph structure underlaying the acitve sources spatial distribution (Figure 5c).

**Figure 5:**
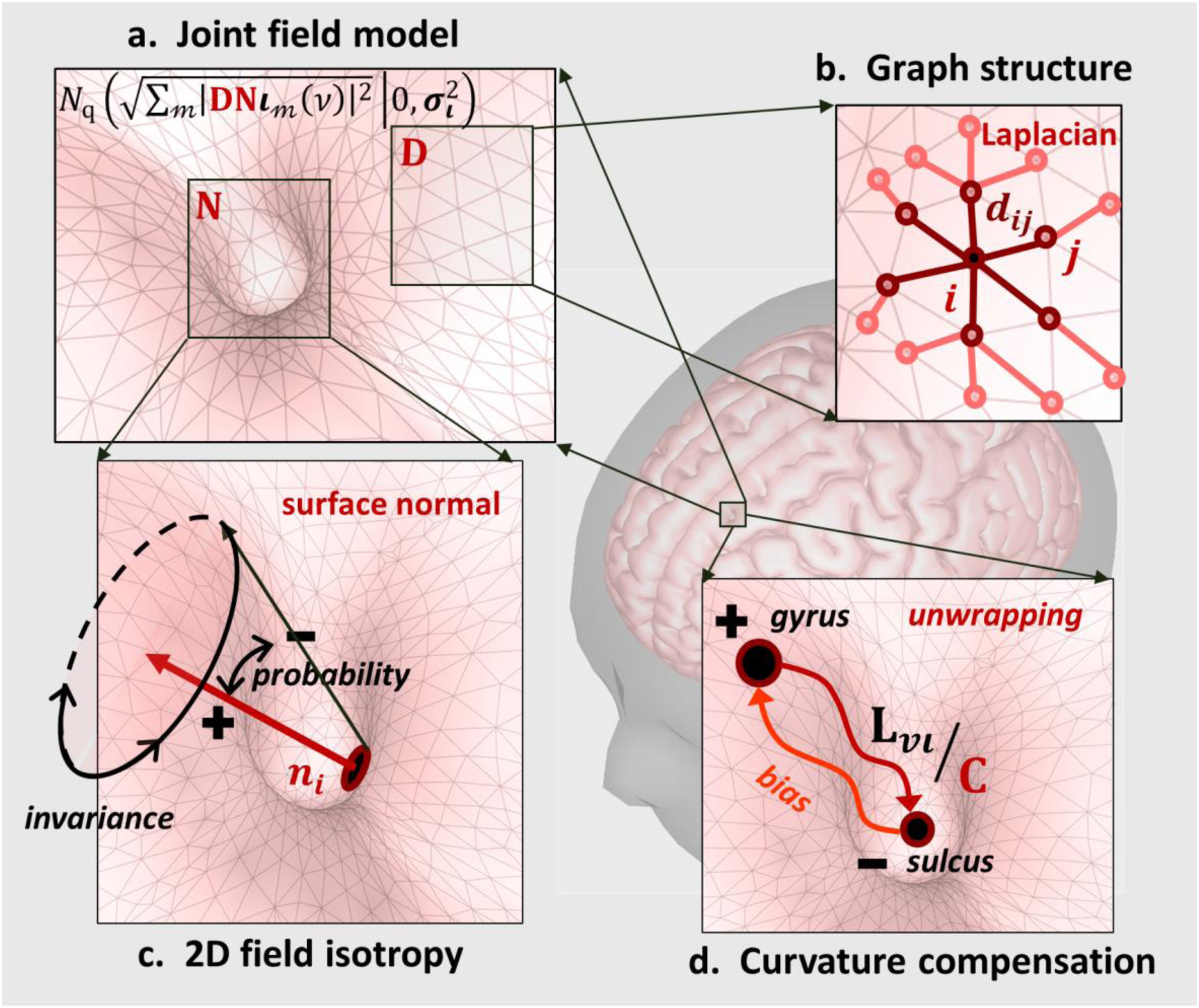
Links introduced in the Gaussian distribution of the spectral activations via the composited effect of two operators. a) first, reducing the field components 3D degeneracy through the normal projection operator (**N**). b) second, introducing dependencies between neighbor activations through the Laplacian operator (**D**). **N** yields rotational invariance of the activations Gaussian probability model, with pivot axis on the surface normal (**n**_**i**_) at every cortical point (**i**). c) the probability decreases as the orientation of activity deviates from the normal direction. Whereas with **D** the probabilities are not assigned individually to cortical points (**i**), but to its linear weighted combination with neighbors (**j**). With weights of the Laplacian matrix elements (**d**_**ij**_). d) compensation of ESI curvature wrap or bias towards gyri. The operator (**C**) represents the curvature of the cortical surface to unwrapping the Lead Field and therefore the ESI solution.

We aknowledge that ESI solutions based on cortical space can be wrapped at or biased towards surface giri due to atenuation of fields in the deeper sulci areas. Therefore we introduce a compensation factor based on curvature coeficients to unwrapping the Lead Field (Figure5d). To avoid the reverse bias towards sulci the final estimator is computed as the average of two preliminar solutions, one favoring the sulci activation and a second one doing so onto the giri.

We also extended the sample-space group penalization to a 3D cartesian space, with the inclusion of two aditional spaces (Figure 6a). First, the space of frequencies within a spectral band carrying on common MEG/EEG scalp topologies (Figure 6b) as typicaly observed in real data. Second, the space of generators within cortical parcels, asuming the coactivation of generators within areas that are defined upon a structural or functional atlas (Figure 6c).

**Figure 6:**
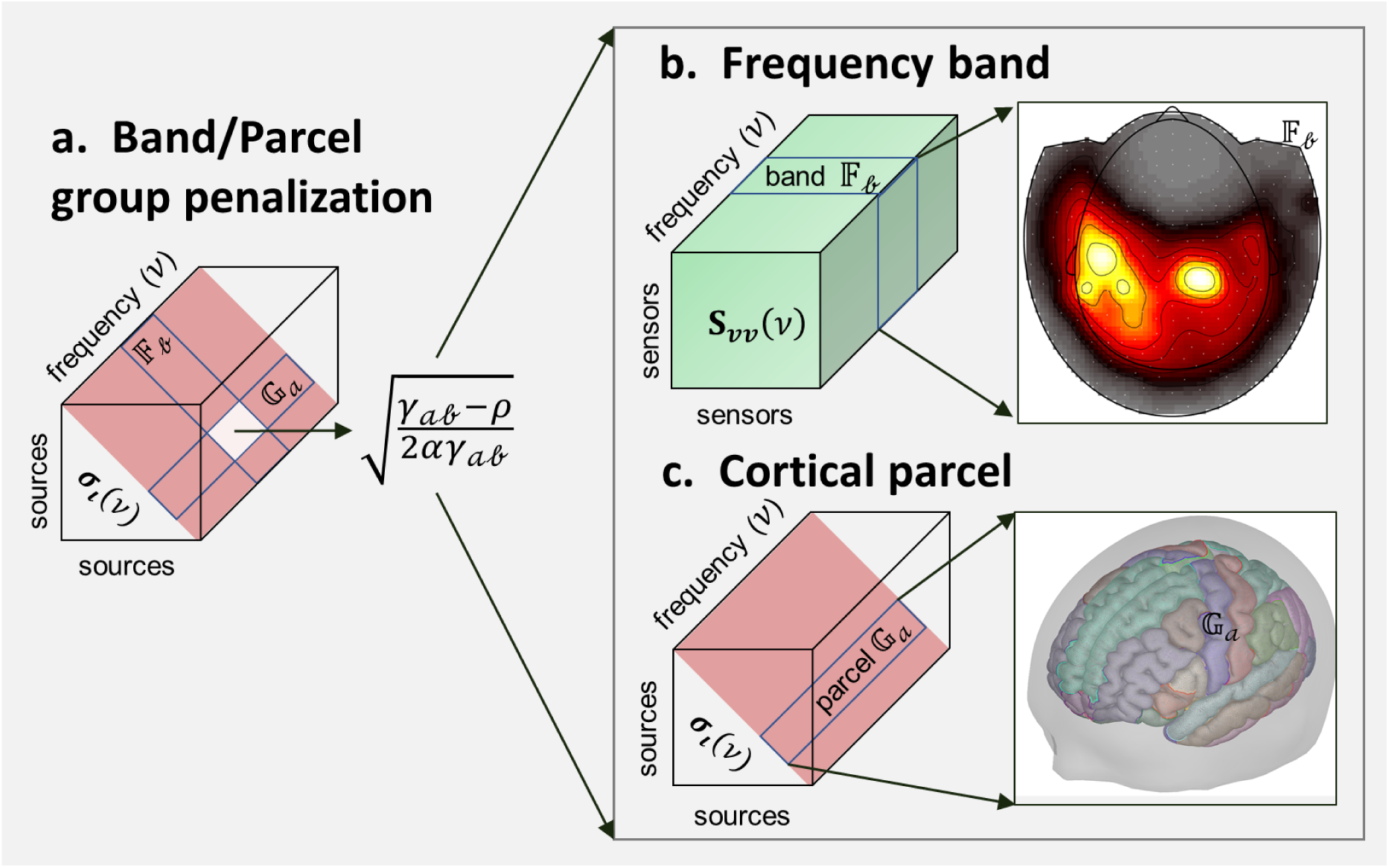
Group penalization of the Hierarchical Elastic Net a) with implications in the estimation of the variances **σ**_**ι**_, which determine the statistical relevance of activation patterns. As a consequence of this penalization the variances are computed from a single parameter γ_𝒶𝒷_ that performs a variable selection of groups of parameters. These are representing certain activations belonging to the interception of two groups 𝔽_𝒷_ (band) and 𝔾_𝒶_ (parcel). b) first, for a spectral range 𝔽_𝒷_ corresponding to an electrophysiological band, with common observable spatial patterns for all the frequency components within this band. c) second, activity corresponding to generators within a cortical area 𝔾_𝒶_, defined into a given neuroanatomical Atlas, which acknowledges activations within delimited areas are highly integrated and thus prone to appear simultaneously.

## Results and Discussion

### Concurrency of sSSBL++ in the low-density pseudo-EEG of a high-density MEG

In evaluating the sSSBL++ performance we diverge from the use of synthetic MEG/EEG signal simulations, typically stablished upon idealization of the spatial distribution and spectral composition of underlaying neural activity [Pascual-Marqui et al 2014; Haufe and Arne 2016]. This is a major flaw of the state-of-the-art validation methods that deserves special attention here. We aim to resolve this by leveraging the concept of concurrency between neuroimaging techniques (Wang, 2019), through introducing a completely new validation benchmark based on the comparison between ESI solutions for the MEG and EEG modalities (Figure 7). One from a high-density real MEG signal (Figure 7a) to establish a landmark for source activation (Figure 7b), that is later used to simulate the pseudo signal of a low-density EEG (Figure 7c). The other, attempting to retrieve the original source activation from the pseudo EEG signal alone (Figure 7d).

**Figure 7:**
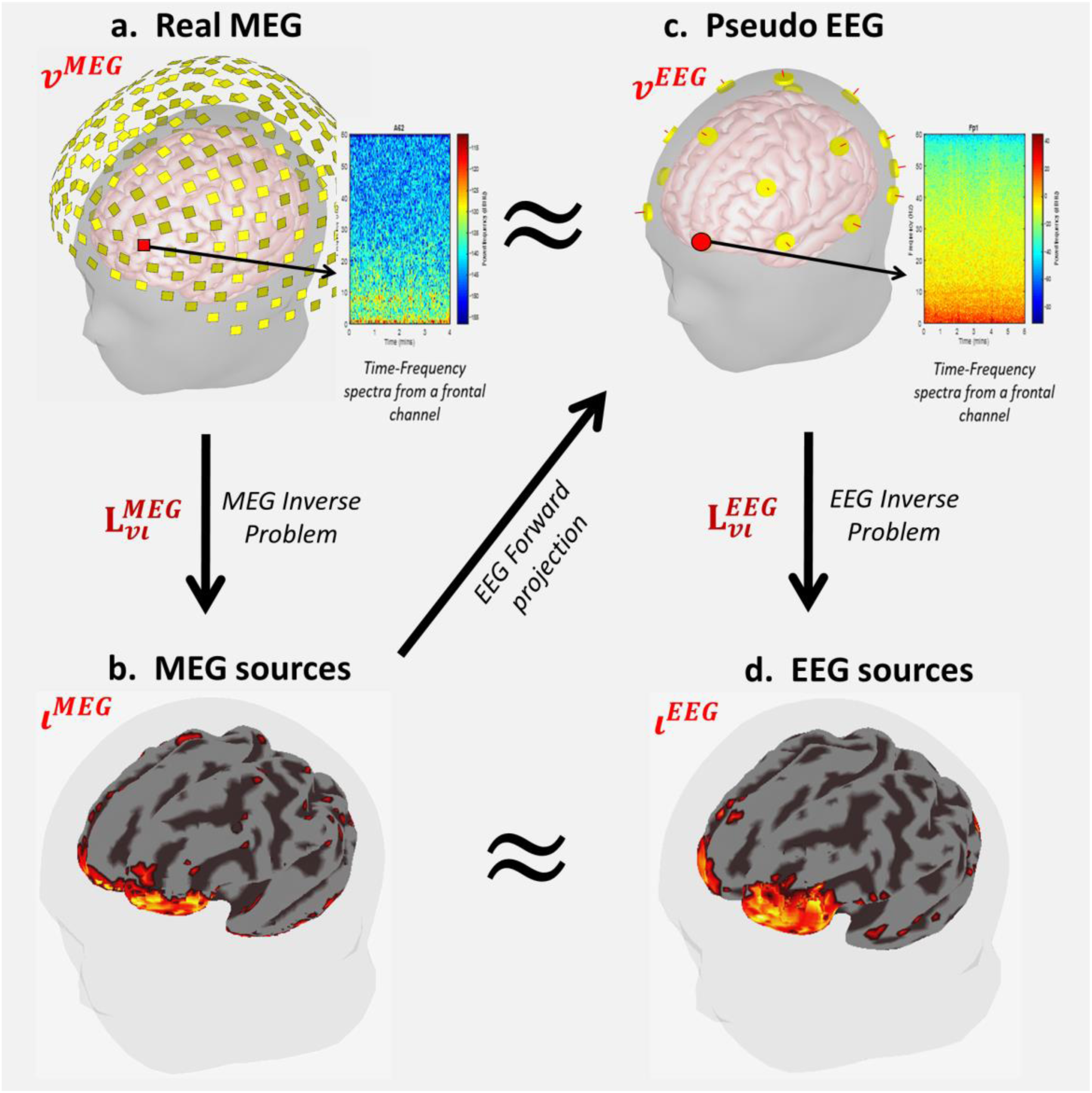
Novel methodology for ESI validation, based on the concurrency between MEG and its pseudo EEG. a) actual distribution of MEG sensors in the subject’s native space and time frequency composition a frontal sensor signal. b) a landmark for alpha band spectral activity construed through sSSBL++ for the MEG signals, with Brainstorm head model of the three overlapping spheres. c) 10-20 EEG system adjusted to the native subject’s space, used to produce pseudo-EEG from the projected MEG alpha source landmark, with a Brainstorm head model of the Boundary Element Method. d) with sSSBL++ we achieve a concurrent pseudo-EEG to judge from the time-frequency composition of the signal for the frontal sensor analogous to that of the MEG. Concurrent alpha source activity retrieved from the pseudo-EEG with the sSSBL++.

Following the scheme of Figure 7 we studied the sSSBL++ concurrency on real MEG -pseudo EEG-data in five bands of frequency (delta 0-4 Hz, theta 4-8 Hz, alpha 8-12 Hz, beta 12-16 Hz and gamma 16-32 Hz). The real MEG was selected from the Human Connectome Project (HCP) database [Van Essen et al 2013], opportunely after its selection by Brainstorm team [Tadel et al 2011] to illustrate the MEG database usage, see details in MM-2.

The sSSBL++ exhibited a good performance in the concurrency test (Figure 8a), to judge roughly for its similarity in performance with typical examples of idealized simulations [Haufe et al 2013, Stokes et al 2017, Van de Steen et al 2019]. In SI-7 we offer an example of the results for sSSBL++ and other methods in such simulations. Meaning that major activations retrieved from the pseudo-EEG across areas coincided with the real-MEG landmark with only minor leakage distortion. This was also supported by the concurrency of fluctuations across frequency from activity in the slow band delta to the faster beta band. For both pseudo-EEG and real-MEG, these transited smoothly from frontal areas (delta) to occipital areas (alpha) as reported by multiple studies [da Silva 2013]. The same effect was observed for the transition bands theta (delta → alpha) and beta (alpha → gamma). Leakage of the pseudo-EEG persisted but relevantly did not overpass specific anatomical areas where major activations were detected.

**Figure 8:**
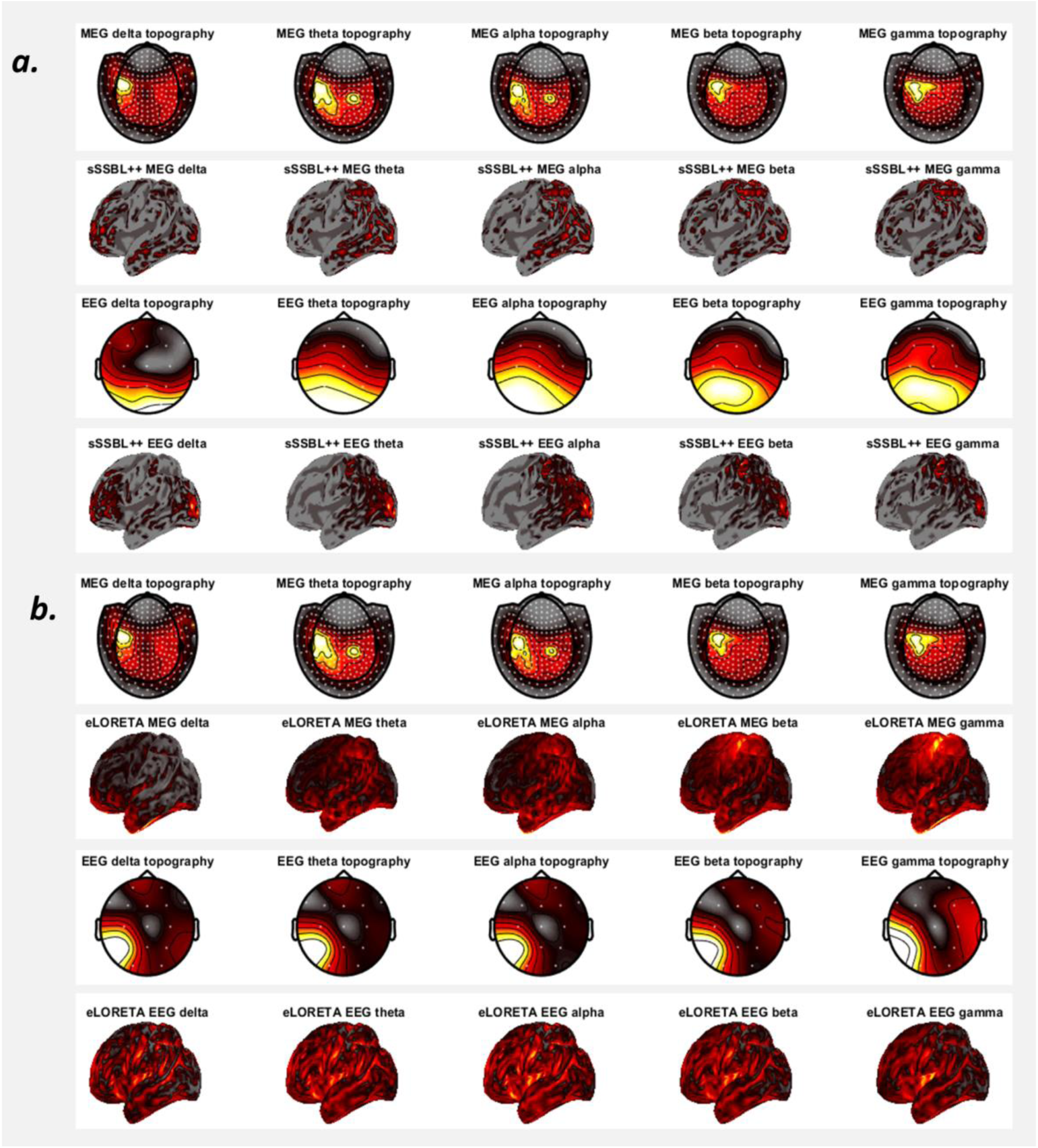
Hot colormaps with results of the concurrency evaluation for a) sSSBL++ and b) eLORETA on 4 spectral bands (delta, theta, alpha, and beta). The topographies at the sensor level represent the cross-spectrum slices for different spectral bands used for the ESI estimation with MEG (top) and EEG (bottom). The performance can be judged qualitative by comparing the source activity (cortical maps) estimated from the MEG (top) and its pseudo-EEG (bottom).

The evaluation was extended to a prevalent instance of ESI: The Exact Low-Resolution Tomographic Analysis (eLORETA) [Pascual-Marqui et al 2011] (Figure 8b). See details on methods used for comparison in MM-2. Roughly, only few activations were expressed concurrently in both the pseudo-EEG and real-MEG eLORETA solutions, which were also persistent across the different frequency bands. Interestingly, their results did defer greatly from those in idealized simulations (Section 7 of SI), meaning in this realistic concurrency scenario its performance was severely affected by leakage.

A quantitative analysis of concurrency, through the surface based Earth Mover’s Distance (EMD) metric [Paz-Linares et al., 2017], confirms this dramatic effect of leakage in eLORETA for all frequencies (Figure9). The EMD quantifies effort to draw leaked activity, produced in the estimation with the pseudo-EEG, back to the MEG landmark. In this situation we use EMD values rated by the ESI differences in the spectral domain, due to the lack of gold standard, in order to discriminate concurrency from bias undergoing ESI estimation, see MM2. The evaluation was extended to the Linearly Constrained Minimum Variance (LCMV), the second generation ESI method selected by HCP for the study of brain connectome based electrophysiology [Van Essen et al 2013; Larson-Prior et al 2013], see MM-2 and SI-2. The leakage for the LCMV solution measured in terms of EMD reaches far beyond that of eLORETA.

**Figure 9:**
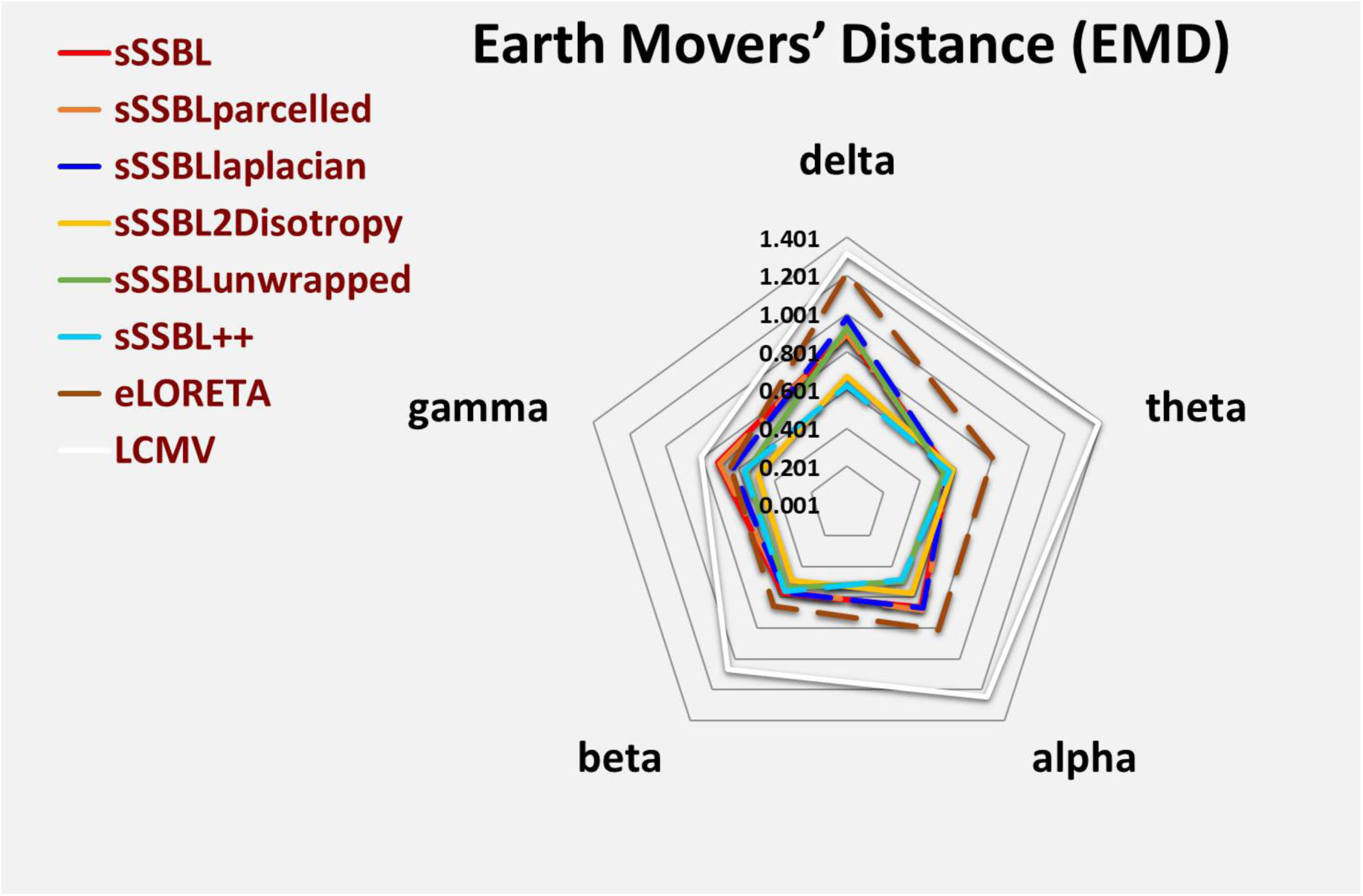
Radar graph with values of Earth Movers’ Distance (EMD) for sSSBL with several instances of priors and state-of-the-art-methods eLORETA and LCMV, in five spectral bands (delta, theta, alpha, beta, and gamma). The priors introduced produce the following effects: spectral smoothness that we refer simply with sSSBL (**Figure 6b**), parcel smoothness sSSBL parcelled (**Figure 6c**), neighbors link sSSBL laplacian (**Figure 5b**), field rotational invariance sSSBL2Disotrpy (**Figure 5c**), curvature compensation sSSBLunwrapped (**Figure 5d**). These were added in isolation to evaluate their effect on leakage independently and altogether denominated sSSBL++. Here the EMD is quantifying the present amount of leakage produced by any method in the concurrency evaluation of **Figure 7**, rated by the EMD for the deviation of these methods across the spectral bands, see MM-2.

The values of EMD for sSSBL++ were representative of the results in Figure 8, lower to those of eLORETA and therefor much lower in comparison to those of LCMV. This quantitative analysis was also extended to several instances of sSSBL++, adding on each of the proposed priors to confirm their possible effect on leakage in isolation. See the summary of these in Figures 5 and 6. Importantly, overall the performance of sSSBL++ was far more superior than that of any instance of sSSBL, only outperformed by the 2D field isotropy in some spectral bands. Also, overall all sSSBL instances performed better than eLORETA and LCMV only falling behind for the gamma band the spectral smoothness (sSSBL) and parcel smoothness (sSSBLparcelled), which performed very similar.

These results suggest, first a universal lack of rigor in ESI validation procedures, which are limited to evaluate the performance only in idealized circumstances [Haufe and Ewald 2016], and second that the effect of leakage in real data for typical ESI methods might be much more severe than usually expected [Van de Steen et al 2019]. Therefore, it would be required for future efforts to consider a validation benchmark like the one proposed here or at least fair simulations of brain activity. Relevantly, in this scenario the sSSBL++ bottom-up approach allows curtailing of this effect considerably.

We acknowledge the concurrency test based on real-MEG/pseudo-EEG, although superior to previous validation methods, is lacking an actual landmark for brain activity. But for different reasons this is not readily accessible by means of noninvasive techniques, e.g. fMRI. Previous works on MEG/fMRI concurrency have provided maps with positively correlated features, but with no account for those expressed exclusively in either technique, which precisely explain their flaws manifested in form of leakage.

### Confirmation of sSSBL++ with the MRI of brain lesions

We propose an alternative solution to discriminate on the effect of leakage in ESI methods based on the comparison of EEG sources with the MRI shine through of hemorrhagic brain lesions. This is possible due to the good definition of this type of lesion in the MRI image and their abnormal electrophysiological responses that could possibly be observed through EEG in any spectral band. In this scenario we define leakage as the responses estimated ESI on the affected areas that could be classified as normal. See details in MM-3.

The MRI and EEG collected belong to an infant born at 36 gestational weeks without any noticeable obstetric or clinical complication (Apgar 9/9, birth weight of 2,370 grams). However, at the age of 2 months an intracranial hemorrhage was detected, caused by hemorrhagic disease of the newborn (HDN). HDN is a common cause of the “acquired homeostatic disorder of newborn infants”, which can take place without the presence of any underlying disorder [Bör et al 2000]. The HDN diagnosis was confirmed through a T2 MRI study, see the three views at Figure10top. This shows clearly an extended subdural hematoma and right hemispheric stroke located in the area surrounding the middle and posterior cerebral artery.

EEG data was recorded a month later (3 months of age) in expectancy of a higher degree of maturity-in terms of electrophysiological brain age- in which not only delta spectral features but the faster band theta may possibly be appreciated [Niedermeyer and da Silva 2005; André et al 2010]. Aiming to reconstruct the source of all neurophysiological activity, that may possibly be expressed in the EEG, we computed the sSSBL++ solution for a broad spectral band (0-7 Hz) unifying the delta and theta bands. See in Figure10 middle the results in red color, where the red emphasis corresponds to activity classified as abnormal for the broad range of electrophysiological frequencies.

**Figure 10:**
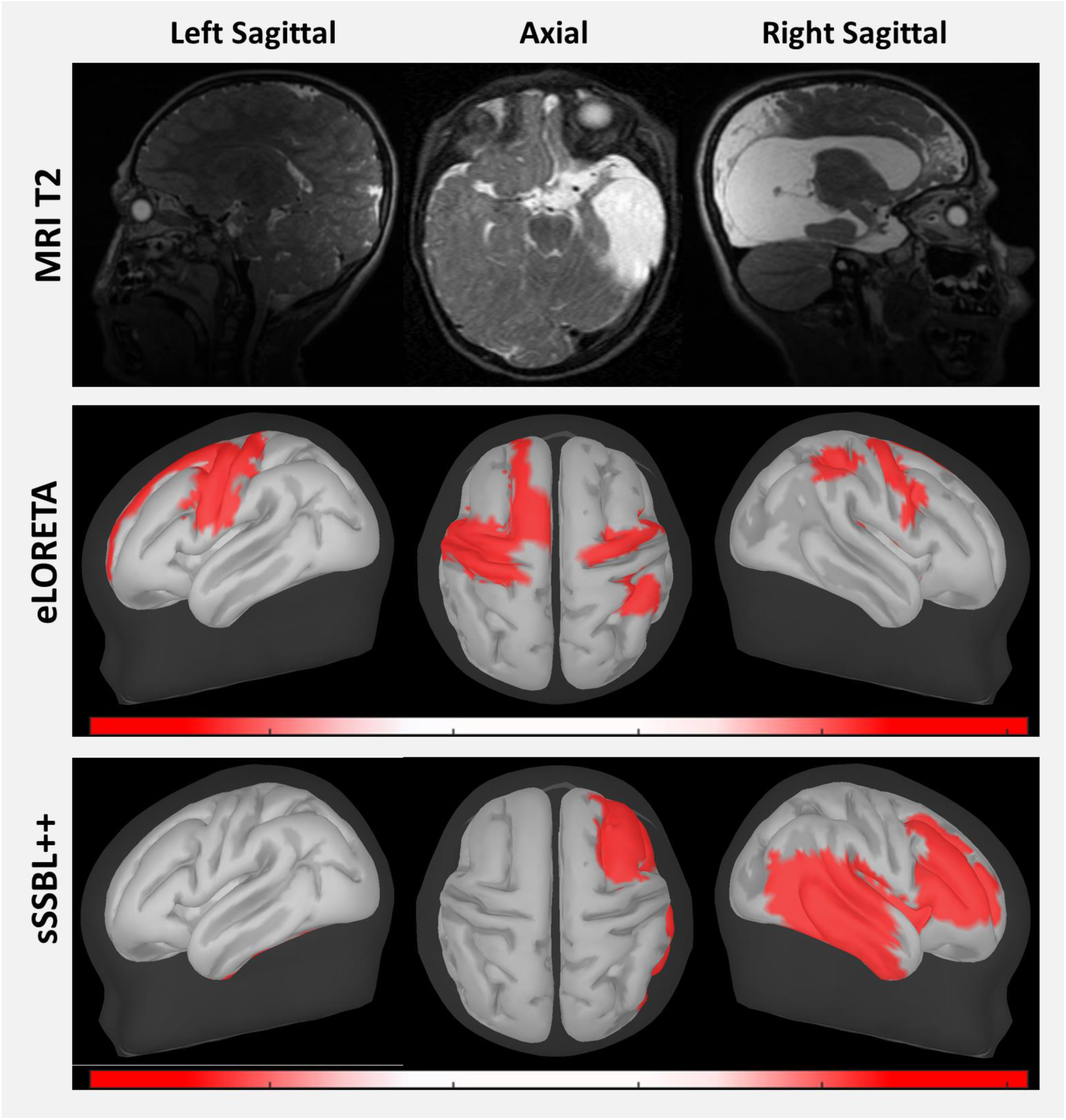
Detection of abnormal source electrophysiological activity caused by hemorrhagic brain lesions that were observed in the MRI T2 of the subject (top). The results of ESI with eLORETA (middle) and sSSBL++ (bottom) were obtained from the signal of a 19 channel EEG in the electrophysiological range of frequencies. The performance can be judged qualitative in terms of the undertone of the areas detected as abnormal by the methods with their MRI T2 shine-through.

These are in high correspondence with the lesions mapped through the T2 MRI, provided all the abnormal areas detected by sSSBL++ were in the left hemisphere. Also, their distribution across different lobes and extension resemble that of the hemorrhage, which involved the occipital, temporal, tempo-parietal, frontal and fronto-parietal areas. Importantly, no perceptible normal activity infiltrating inwards to affected areas could be observed in the sSSBL++ solution. Rather, the areas detected as abnormal were extended far beyond the lesions.

On the contrary the abnormal activity detected with eLORETA exhibited great differences with the MRI shine through of the lesions. Most of the abnormal activity was detected in the left hemisphere which cannot be justified to judge for the normal T2 contrast of this. Also, in the right hemisphere there was a wide mismatch between the shine through of lesions and the abnormal areas detected by eLORETA.

Notice that the ESI solutions were computed with an average head model and therefore in different space as the MRI. A T1 MRI -adequate structural space to define the individual head model- was not registered anticipating the typically deficient tissue contrast observed in T1 relaxation of newborns. An adequate quantification of the present amount of leakage would require projection of both the T2 MRI and EEG to a common structural space. Up to now we lacked a processing pipeline that could perform with acceptable precision the registration, segmentation and head model construction based on the T2 MRI of infants. This process, if performed by means of the standardized tools, would produce a roughly approximated head model, therefore requiring further developments.

## Materials and Methods (MM)

### MM-1 sSSBL algorithm

sSSBL pursues estimation of the cross-spectrum through a maximum “evidence” search via the Expectation-Maximization algorithm. Where evidence is defined as the conditional probabilities of two groups of parameters, given the available data samples or MEG/EEG cross-spectrum **S**_***vv***_(*v*): these are *σ*_***ι***_, that controls the statistical relevance the source cross-spectral components **S**_***ιι***_(*v*), and *σ*_***ξ***_, that controls the level of noise of the observations. This is done through an iterated scheme that produces an approximated representation of the evidence (expectation) followed by its maximization, that guarantees convergence to a local maximum. See in Figure 11 the algorithm for the expectation step illustrated in graphical form. In SI-5 and SI-6 we provide the derivation of the later, extended with all priors included in the sSSBL++.

**Figure 11:**
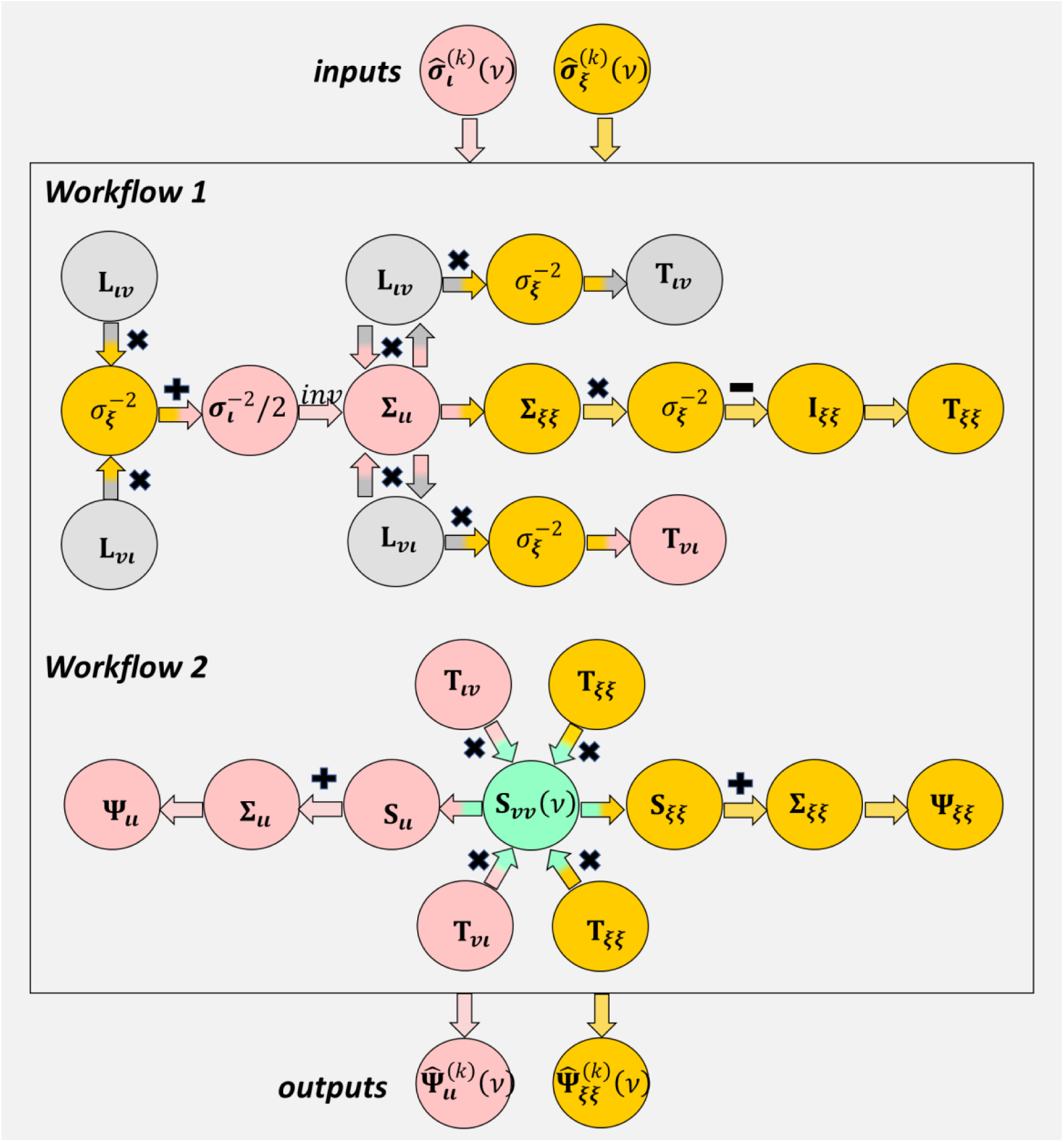
Graphical representation of the sSSBL Expectation algorithm that precedes the Maximization. The red color identifies quantities (circles) and mathematical operations (arrows) associated to source activity, whereas the green identifies those for observation noise. The constant **L**_**vι**_ (Lead Field) and its transconjugated **L**_**ιv**_ is identified with a gray circle. Workflow 1 uses the variances of noise 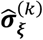 and activations 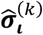 to produce two locally linear transfer operators: one asymmetric for the re-estimation of ESI solution (**T**_**ιv**_ and its transconjugated **T**_**vι**_) and another one symmetric for the noise re-estimation **T**_**ξξ**_. Workflow 2 produces the effective empirical covariances of source activity 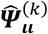 and noise 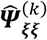, used in the Maximization step.

The maximization step is carried out via estimation formulas that we refer to here as the group Hierarchical Elastic Net, see Figure 12. This resembles the estimation formulas of the vector regression Elastic Net [Paz-Linares et al., 2017] and the Sparse Bayesian Learning [Wipf et al., 2009], but in this case through the arithmetic mean ***ψ***_***ι***_ of typical vector regression inputs corresponding to the samples. See the derivation of these formulas in SI-7 also extended to the sSSBL++. Although not shown here, we perform control of the global sparsity level through estimating the parameters *α* and *ρ*, in completely analogous form to the procedure in [Paz-Linares et al., 2017], see SI-7.

**Figure 12:**
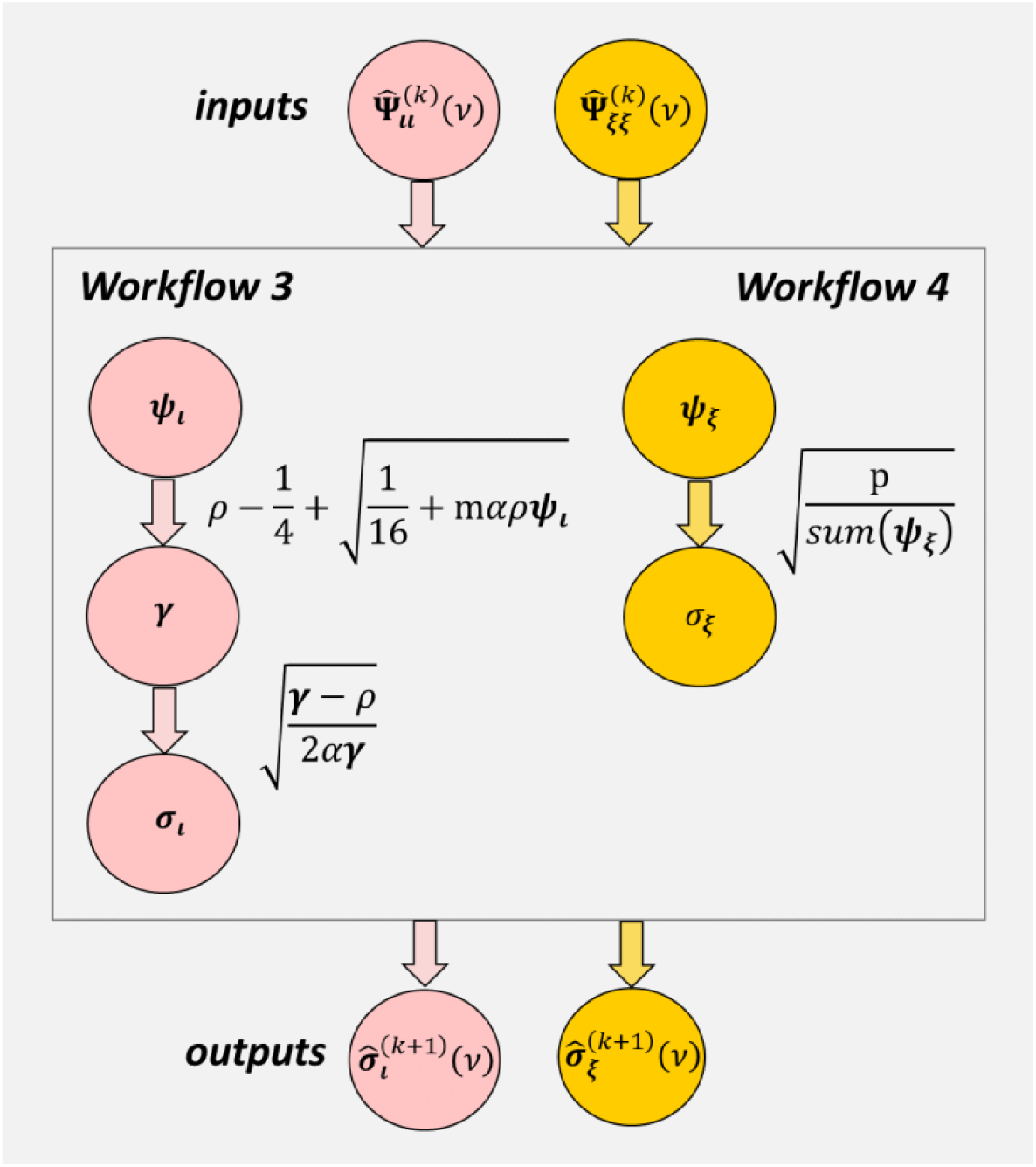
Graphical representation of the sSSBL Maximization algorithm, following the Expectation. The red color identifies quantities (circles) and mathematical operations (arrows) associated to source activity, whereas the green identifies those for observation noise. Workflow 3 uses the effective empirical covariance of the activations 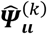 to produce the source variance 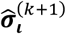. Workflow 4 uses the effective empirical covariance of the noise 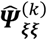 to produce the noise variances 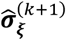. The variances are used in the Expectation step of the next iteration.

To help with the reproducibility we provide in SI-8 a standard pseudocode (nongraphical) and all formulas in a compact way. The code is freely available in GitHub as part of a more general toolbox for MEG/EEG source analysis (<u>BC-VARETA</u>) [Paz-Linares et al 2018; Gonzalez-Moreira et al 2018a; 2018b; 2018c].

### MM-2 MEG/EEG concurrency study

The MEG corresponded to 246 channel preprocessed resting state data of the subject 175237 from the HCP database [Van Essen et al 2013; Larson-Prior et al 2013]. We construed the MEG head model following the Brainstorm pipeline [Tadel et al 2011], designed specifically to utilize HCP structural and functional data, and which illustrates the head model processing with the subject cited above (175237). The cross-spectral tensor used in ESI (Figure3a3) was computed at a frequency resolution of about 0.5 Hz from 172 Fourier coefficients samples (trials), extracted through the Fourier transformation of the recordings spanning about 2 seconds at a sampling rate of 508Hz for each trial. The EEG head model corresponded to 19 channels in the 10-20 system, computed also following Brainstorm recommendations via Boundary Element Method upon 3 tissue layers (inner skull, outer skull, scalp) extracted with FSL [Smith et al 2004; Jenkinson et al 2012] and on 6K generators of a discretization of the midthickness.

We performed the concurrency evaluation, according to the illustration in Figure8, following three steps. First, the inversion of real MEG spectral data upon the MEG head model, via ESI with a given method. Second, the generation through the EEG head model of the pseudo EEG spectral data, corresponding to spectral activity determined in the first step. Third, the inversion of the pseudo EEG spectral data upon the EEG head model, via ESI with the same method as in the first step. The code and data to reproduce these results is freely available in GitHub (<u>sSSBL-Concurrency</u>).

One of the two methods proposed for comparison (eLORETA) belongs to the class of linear methods therefore not provided of any capacity towards bottom-up leakage control (see SI-2) [Pascual-Marqui et al., 2006]. The other method (LCMV) was a Beamformer that belongs to the second ESI generation, which can provide bottom-up control of leakage. But it does partially, due to its lack of a specific spectral model. Nevertheless, we leverage a more efficient implementation provided in fieldtrip that provides eLORETA the capacity to perform the direct computation of the source cross-spectrum via a linear transfer function and with regularization parameters that can be adjusted with complex variable cross-validation [Oostenveld et al. 2011].

To judge the quality of methods or leakage in the concurrency evaluation we used an implementation of Earth Movers’ Distance on surface [Paz-Linares et al., 2017]. In this case their values for the difference between the MEG and EEG solutions in the concurrency test were rated by their values for the difference between delta and alpha bands of the MEG. This turns the validation metrics sensitive to detecting possible bias of the methods, towards certain cortical areas, as with the case of eLORETA and LCMV. Bias makes it difficult to judge, in the absence of the “gold standard” as is the case of our concurrency evaluation, whether the present similarities can be explained with good quality or with low sensitivity to changes in the spectral domain.

### MM-3 MRI/EEG confirmatory study

Acquisitions like the EEG and T2 MRIs used in the confirmatory study here are part of the routine examination following the admission of cases in the neurotherapeutic area of the Neurodevelopment Research Unit located in Queretaro campus of UNAM. The data used for the confirmatory study will be available upon request of the readers.

EEG was recorded in resting state condition with eyes closed on 19 channels defined in the 10-20 system using a Medicid™ IV (*Neuronic Mexicana, S.A.; Mexico*). The contact impedance for all electrodes was tuned down to a level of 5 kΩ or below and the bandwidth of the amplifiers was set within the limits of 0.3 Hz to 30 Hz. The preprocessing was performed off-line and through manual artifact rejection guided by expert hands on the standards of the International Federation of Clinical Neurophysiology (IFCN). The preprocessing yielded an amount of 16 artifact-free trails each spanning over 2.56 seconds. To construct the head model for ESI we used the Brainstorm default anatomy ICBM152, via Boundary Element Method upon 3 tissue layers (inner skull, outer skull, scalp) and on 6K generators of a discretization of the midthickness.

Structural T2 MRIs were acquired using a 3T scanner (General Electric Healthcare, Milwaukee, Wisconsin, US) with 16-channels of a head neurovascular coil (HDNV). The infant was awake, laying back and wearing earplugs for protection against the MR room noises. Figure11c shows structural images acquired with two different pulse sequences: A T2-weigthed spin echo sequence (Left and Right), TR/TE 2500/68 ms, flip angle 90°, slices 196, slice thickness 1 mm, matrix 224 × 224, FoV 220 × 220 mm2, voxel sizes of 0.8 × 1.0 × 0.8 mm3. A T2-weigthed turbo spin echo sequence (Center), TR/TE 7000/100 ms, flip angle 111°, slices 70, slice thickness 2 mm, matrix 448 × 352, FoV 220 × 220 mm2, voxel sizes of 0.4 × 0.4 × 2.0 mm3.

The classification of abnormal activity was carried out in statistical comparison with norms for EEG source activity in infants of the same range of age [Otero et al 2011]. For each method (eLORETA and sSSBL) the norm of source activity was computed using the EEG of 179 newborns collected on the basis discussed at the beginning of this section. Then, the estimated activity for the pathological patient for every method was compared to the mean and standard deviation of its norm. The criteria for the selection of abnormal areas was according to the difference, across all frequencies, between the norm mean and the estimated activity for the patient. The threshold used for classification was twice the standard deviation of the norm.

## Conclusions

We demonstrated that the impact of leakage in common Electrophysiological Source Imaging methods can be much more severe than expected. Shedding light on the actual extent of leakage required completelly new validation procecedures slanted towards a landmark based on the real data rather than in simulations of Brian electrical activity. This was furnished here via two different instances, that according to our knowledge are lacking of any precedent and may else serve to future efforts: First, concurrency of the ESI solution in the low-density pseudo-EEG of a high-denstity MEG. Second, comparison against the MRI shine through of brain lesions which turn the affected areas into a source of abnormal electrophysiological activity.

We proposed ESI via direct sparse Bayesian learning of spectral responses -coined spectral Structured Sparse Bayesian Learning (sSSBL) and its extension with multiple priors (sSSBL++). This procedure is hitched to technical difficulties for the statistical learning of complex valued data models, that we have solved here. The results with sSSBL++ display far more control of leakage benchmarked in our validation framework against commonly used methods eLORETA and LCMV. This strongly supports our working hypothesis of bottom-up control on leakage, where a possible suggested route is through using ESI priors and learning process directly to the ultimate estimators targeted by leakage.

## Acknowledgments

This study was supported by Grant 61673090 from the National Nature Science Foundation of China, and Grants DGAPA-IN200917, CONACYT 4971 and CONACYT 251309 from the National Council on Science and Technology of Mexico.

## Contributions

EGM and DPL designed the method and wrote the paper. AAG, YW and ML performed validation and implementation of software platforms. JBB guided MEG/EEG analysis and confirmation of the method. TH senior author who designed the experimental confirmation of the method and collected the data. PAVS senior author who introduced the theoretical background.

## Supporting Information (SI)

### SI-1 Mathematical notation

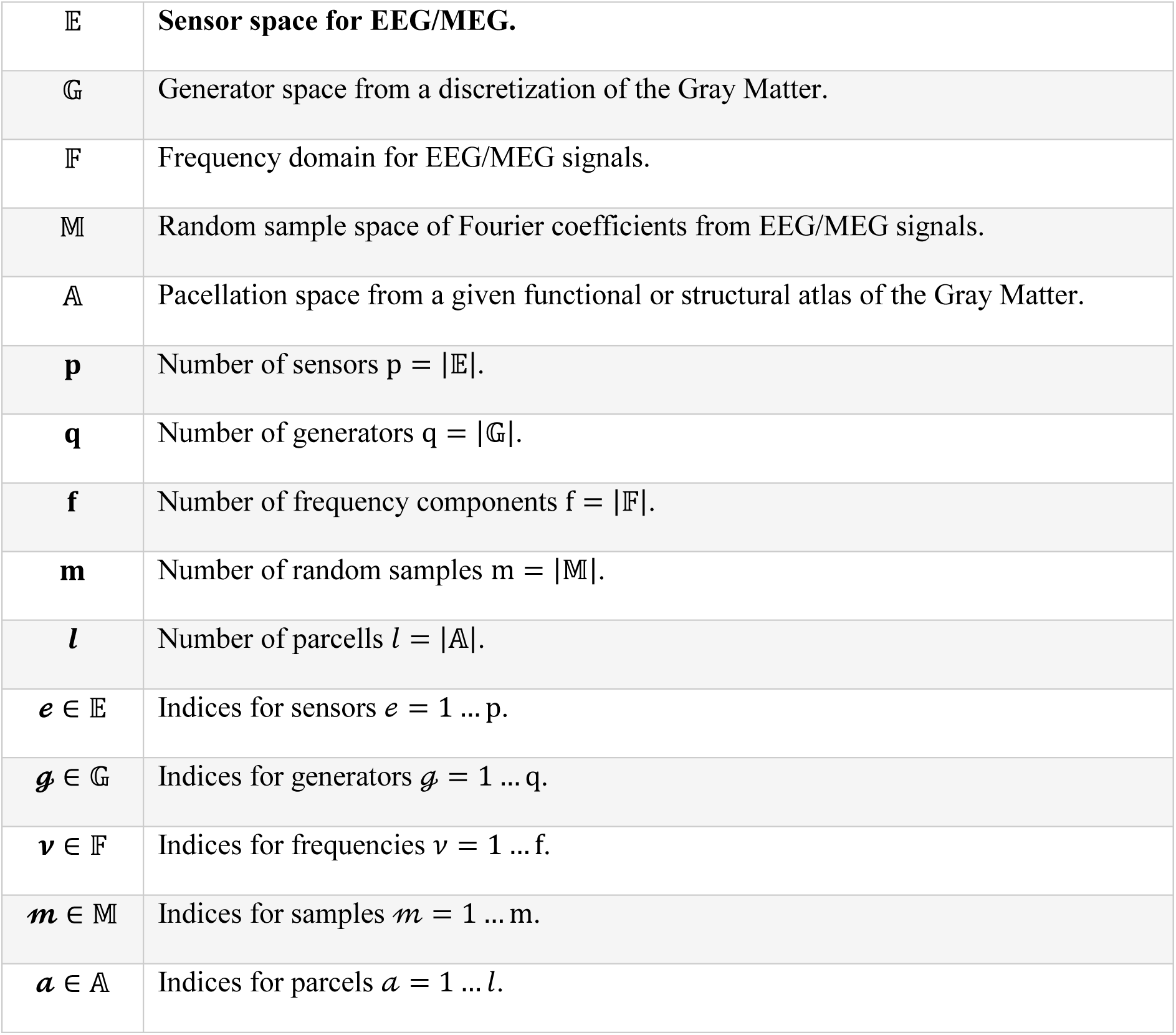
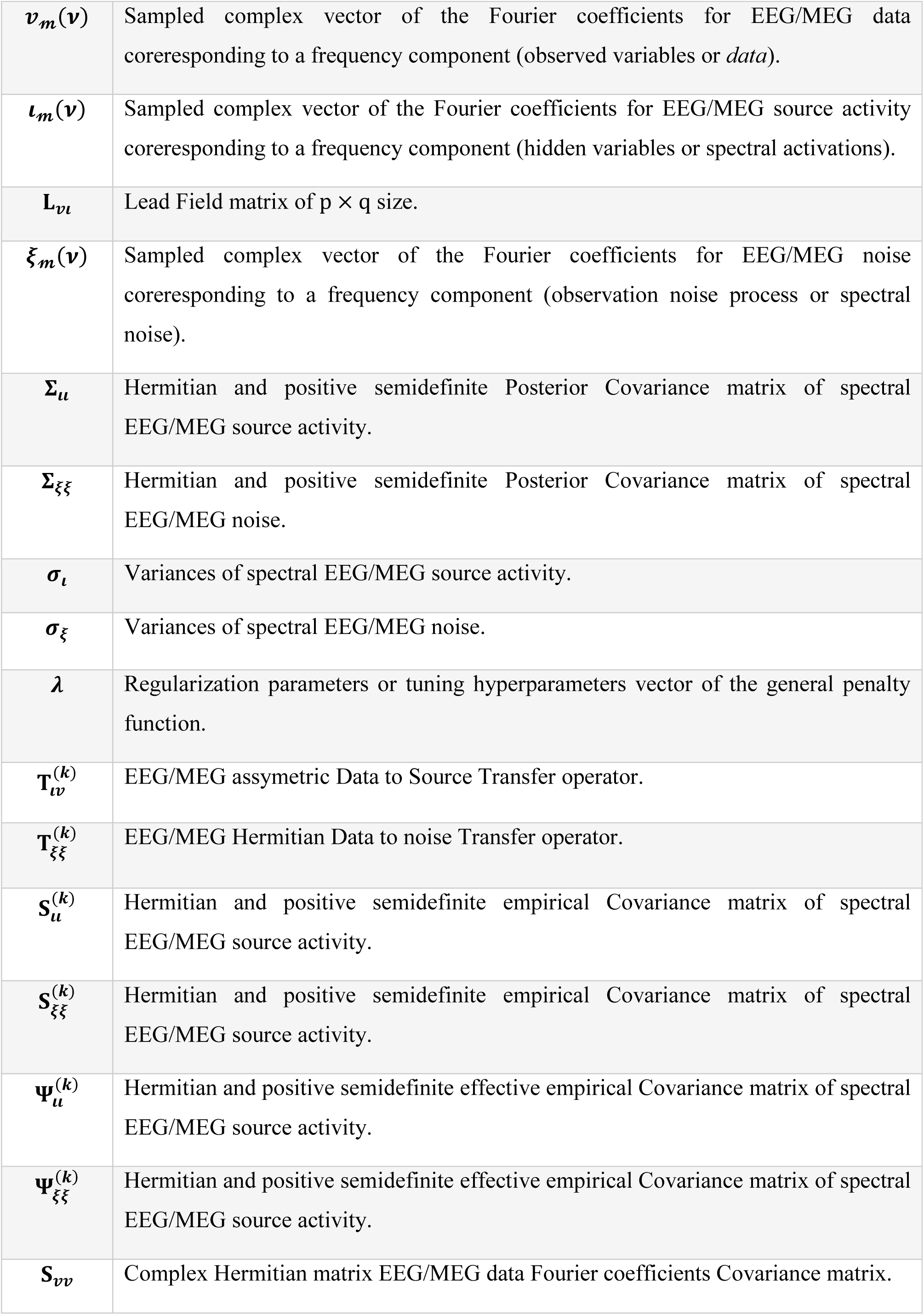

|  |  |
| --- | --- |
| $\mathbb{E}$ | **Sensor space for EEG/MEG.** |
| $\mathbb{G}$ | Generator space from a discretization of the Gray Matter. |
| $\mathbb{F}$ | Frequency domain for EEG/MEG signals. |
| $\mathbb{M}$ | Random sample space of Fourier coefficients from EEG/MEG signals. |
| $\mathbb{A}$ | Pacellation space from a given functional or structural atlas of the Gray Matter. |
| $\mathbf{p}$ | Number of sensors $p = \vert \mathbb{E}\vert $. |
| $\mathbf{q}$ | Number of generators $q = \vert \mathbb{G}\vert $. |
| $\mathbf{f}$ | Number of frequency components $f = \vert \mathbb{F}\vert $. |
| $\mathbf{m}$ | Number of random samples $m = \vert \mathbb{M}\vert $. |
| $\mathbf{l}$ | Number of parcells $l = \vert \mathbb{A}\vert $. |
| $\mathbf{e} \in \mathbb{E}$ | Indices for sensors $e = 1 \dots p$. |
| $\mathbf{g} \in \mathbb{G}$ | Indices for generators $g = 1 \dots q$. |
| $\mathbf{v} \in \mathbb{F}$ | Indices for frequencies $v = 1 \dots f$. |
| $\mathbf{m} \in \mathbb{M}$ | Indices for samples $m = 1 \dots m$. |
| $\mathbf{a} \in \mathbb{A}$ | Indices for parcells $a = 1 \dots l$. |
| $\mathbf{v}_m(\mathbf{v})$ | Sampled complex vector of the Fourier coefficients for EEG/MEG data coreresponding to a frequency component (observed variables or *data*). |
| $\mathbf{l}_m(\mathbf{v})$ | Sampled complex vector of the Fourier coefficients for EEG/MEG source activity coreresponding to a frequency component (hidden variables or spectral activations). |
| $\mathbf{L}_{vl}$ | Lead Field matrix of $p \times q$ size. |
| $\xi_m(\mathbf{v})$ | Sampled complex vector of the Fourier coefficients for EEG/MEG noise coreresponding to a frequency component (observation noise process or spectral noise). |
| $\Sigma_u$ | Hermitian and positive semidefinite Posterior Covariance matrix of spectral EEG/MEG source activity. |
| $\Sigma_{\xi\xi}$ | Hermitian and positive semidefinite Posterior Covariance matrix of spectral EEG/MEG noise. |
| $\sigma_l$ | Variances of spectral EEG/MEG source activity. |
| $\sigma_\xi$ | Variances of spectral EEG/MEG noise. |
| $\lambda$ | Regularization parameters or tuning hyperparameters vector of the general penalty function. |
| $\mathbf{T}_{uv}^{(k)}$ | EEG/MEG assymetric Data to Source Transfer operator. |
| $\mathbf{T}_{\xi\xi}^{(k)}$ | EEG/MEG Hermitian Data to noise Transfer operator. |
| $\mathbf{S}_u^{(k)}$ | Hermitian and positive semidefinite empirical Covariance matrix of spectral EEG/MEG source activity. |
| $\mathbf{S}_{\xi\xi}^{(k)}$ | Hermitian and positive semidefinite empirical Covariance matrix of spectral EEG/MEG source activity. |
| $\Psi_u^{(k)}$ | Hermitian and positive semidefinite effective empirical Covariance matrix of spectral EEG/MEG source activity. |
| $\Psi_{\xi\xi}^{(k)}$ | Hermitian and positive semidefinite effective empirical Covariance matrix of spectral EEG/MEG source activity. |
| $\mathbf{S}_{vv}$ | Complex Hermitian matrix EEG/MEG data Fourier coefficients Covariance matrix. |

### SI-2 ESI concepts and generations

The derived inverse problem of ESI contains three main elements: (i) EEG/MEG data observed in p-sensors; (ii) anatomically constrained EEG/MEG space of q-generators from a brain tissue segmentation of cortical mid-thickness surface or volume; (iii) px3q Lead Field matrix (an electromagnetic vector field weighting the contribution q generators) given a volume conductor model that entails nonbrain tissue segmentation, sensor positioning and nature of observed data (electric or magnetic fields) [Grech et al 2008].

Estimation of the neural causes can be done by inversion of the EEG forward model [Hämäläinen et al 1994; Pascual-Marqui et al 1994; Pascual-Marqui 1999]. In this case we propose a space-frequency analysis by directly inverting its spectral equivalent in virtue of the linearity of the Discrete Fourier Transform (DFT):

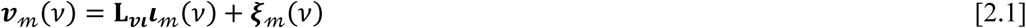

where *v* represents frequencies in the spectral domain, 𝓂 time segments used to compute the DFT, **L**_***vι***_ is the p × 3q Lead Field matrix. The spectral vectors in the equation represent the EEG ***v***_𝓂_(*v*) of size p × 1, source activation ***ι***_𝓂_(*v*) of size 3q × 1 and a sensor noise process ***ξ***_𝓂_(*v*) of size p × 1

Usually the vector fields (**L**_***vι***_(*i*, :) and ***ι***) are projected by the scalar product with another vector field ***d*** (structural not electromagnetic) of Gray Matter tissue directions yielding only q-p degeneracy (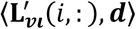 and ⟨***ι, d***⟩).

ESI methods target solving the 3q-p degeneracy EEG/MEG inverse problem kernel (space defining all possible solutions that could explain the recordings) in analogy to the general inverse problem [Groetsch et al 1993]. Reducing the degeneracy beyond further anatomical constraints (Gray Matter ROIs or tissue directionality) is only achievable through regularization or inclusion of some mathematical a priori information [Hämäläinen et al 1994, Baillet et al 1997, Friston et al 2008], which produces a new system with stable unique solution.

The pitfall of leakage has been followed by the deployment of several “generations” of ESI methods within just two decades [Gonzalez-Moreira et al 2018b]. The term “generation” coins basic differences of the statistical learning method towards the construction of the EEG/MEG source transfer operator -via Linear, Nonlinear Univariate, Nonlinear Multivariate models- and the inclusion of prior information in order to ameliorate Leakage. See in Table1 a deeper reconnoitering of such models.

**Table 1:**
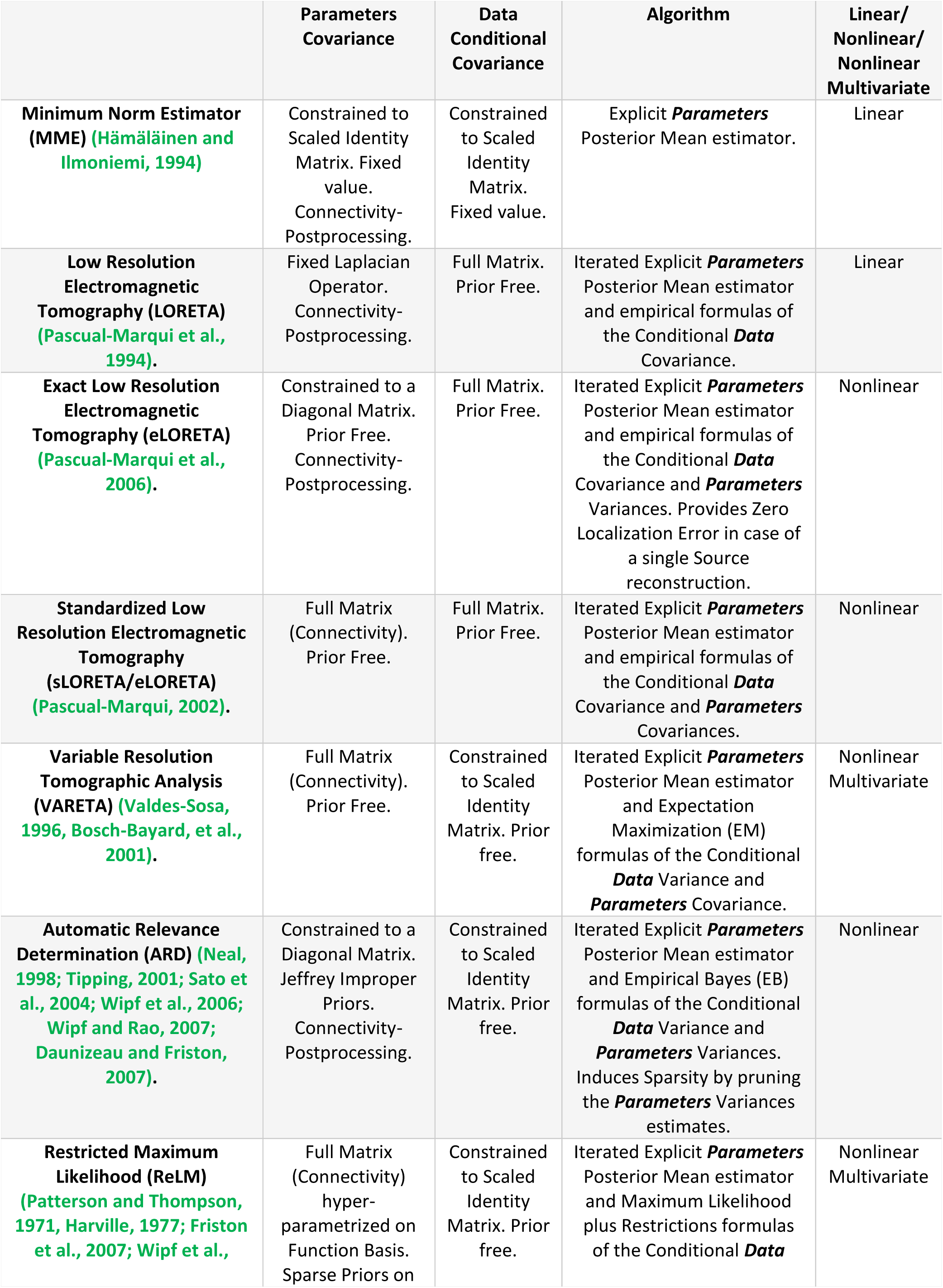
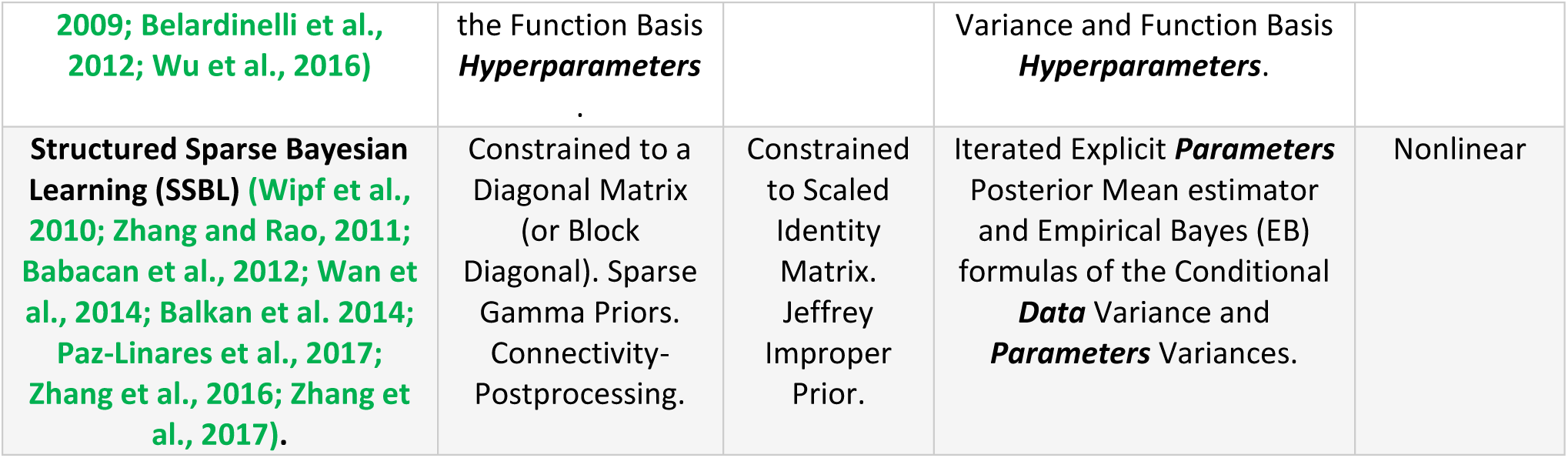
Family of Linear/Nonlinear univariate/Nonlinear multivariate ESI methods.

Summarizing ESI methodologies, we may refer as **Zero-generation** those endeavors of the pre-ESI period, pledged with the search of association between sensor level information and processes at the Gray Matter level [Blinowska et al 2004, Kus et al 2004, Blinowska 2011, Sakkalis 2011], and which has been fairly haunted by critics [Haufe et al 2013, Stokes et al 2017, Van de Steen et al 2019].

**First-generation**, linear nature ESI solution which imposes regularization on the LF ill-conditioning via static priors and which reflects -due it independence of the sensed activity and linearity-similar properties of the Zero-generation with exception of improved anatomical association, e.g. Minimum-Norm Estimates (MNE) [Hämäläinen et al 1994] and LORETAs [Pascual-Marqui et al 1994, Pascual-Marqui 1999].

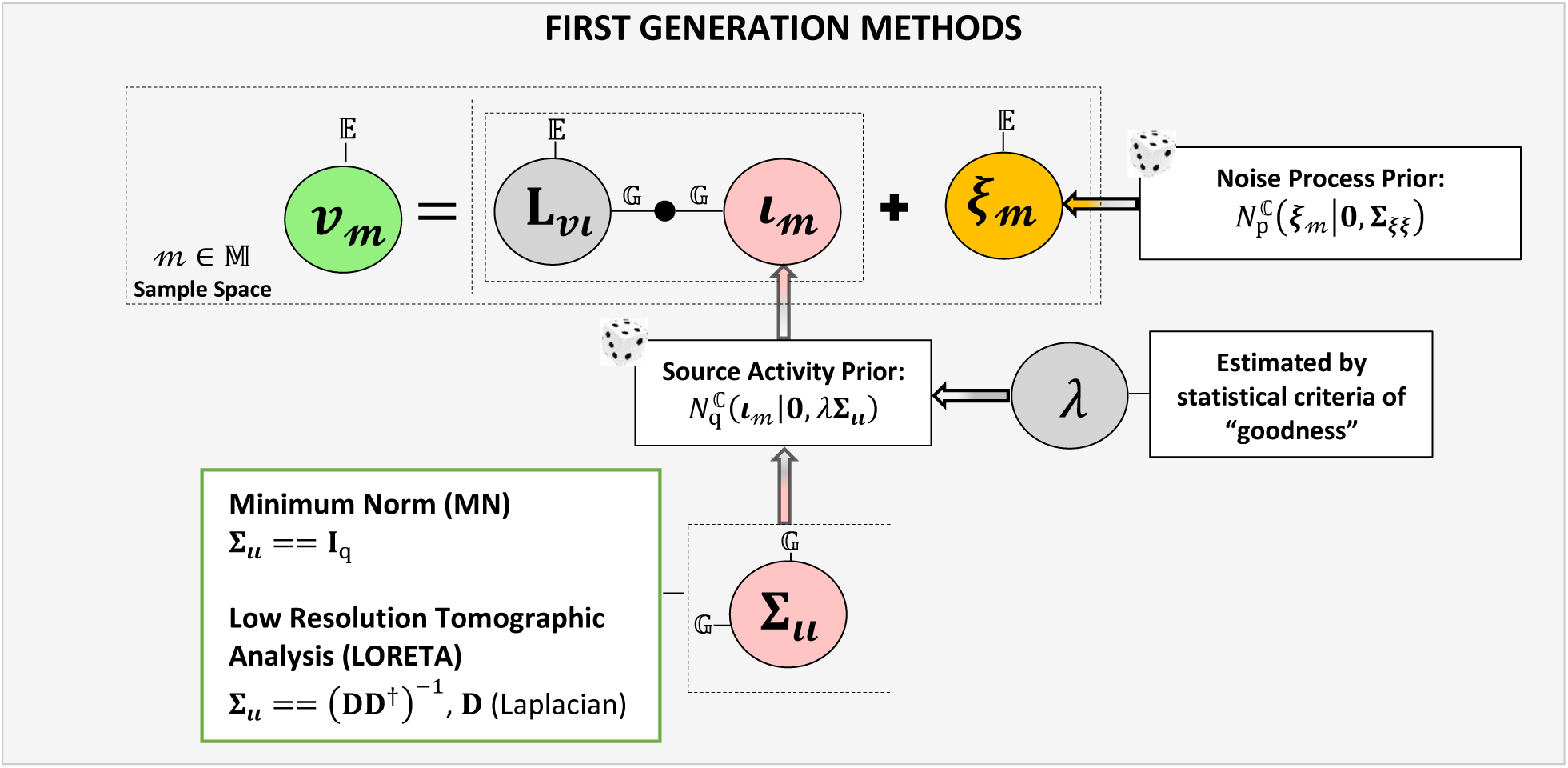

**Second-generation**, nonlinear nature ESI solution for which the transfer function depends on the sensed activity and that usually pursues a sparsity pattern of activation, introduced in form of flexible and univariate priors that improve variable selection, e.g. Exact LORETA (eLORETA) [Pascual-Marqui et al 2006, Pascual-Marqui, 2007], Multiple Penalized Least Squares (MPLS) [Vega-Hernández et al., 2008], and Structured Sparse Bayesian Learning (SSBL) [Paz-Linares et al., 2017].

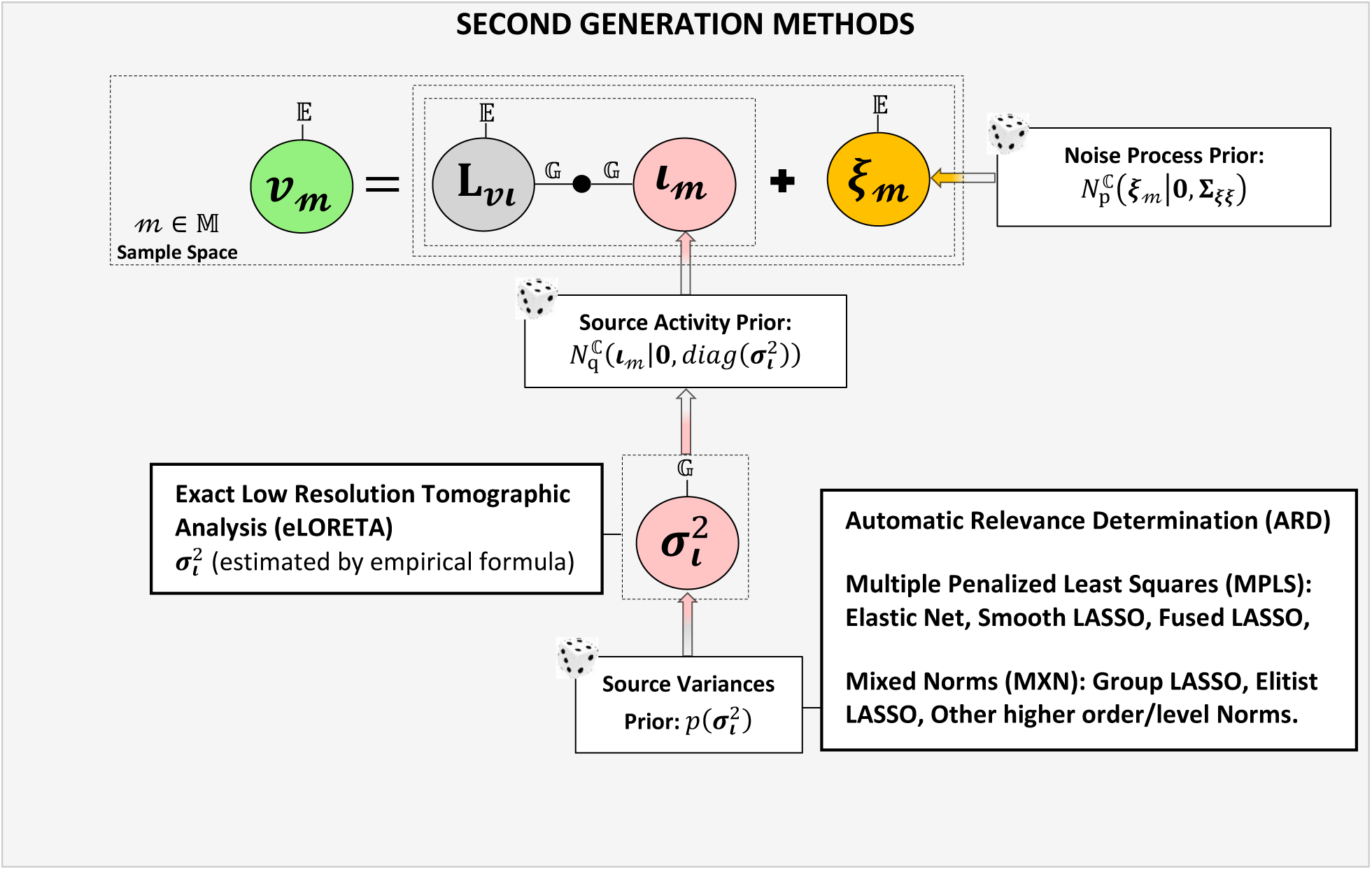

**Third-generation**: nonlinear and multivariate nature ESI solution which acknowledges possible coactivation due to the link between active sources, and therefore leads in some cases to improved determination of activity and its statistical dependencies, e.g. Variable Resolution Tomographic Analysis (VARETA) [Valdes-Sosa et al 1996, Bosch-Bayard et al 2001], Restricted Likelihood Maximization (ReLM) [Patterson and Thompson 1971, Harville 1977, Mattout et al 2006, Friston et al 2007, Friston et al 2008, Wipf et al 2009, Belardinelli et al, 2012] and Brain Connectivity VARETA (BC-VARETA) [Paz-Linares et al 2018, Gonzalez-Moreira et al 2018a, 2018b, 2018c]. *(ESI generations)*

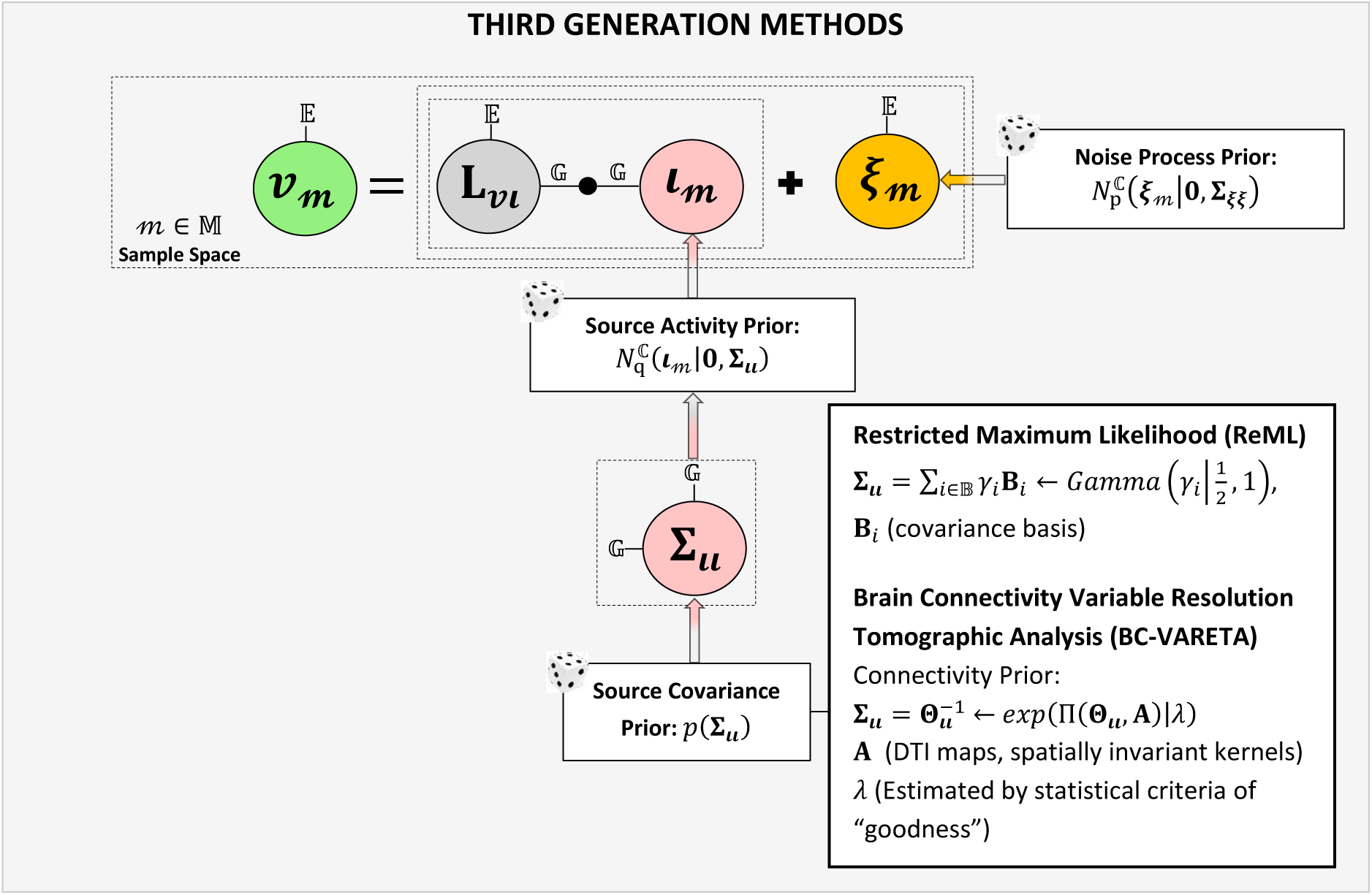

### SI-3 Andrews and Mallows Lemma for the complex variable Hierarchical Elastic Net

The results hierarchical Elastic Net can be extended to complex domain by modifying the corollary of Andrews and Mallows Lemma for Real ENET Gibbs pdf (Gaussian-Laplace) in [Andrews and Mallows, 1974]. For the complex case, the integral representation holds.

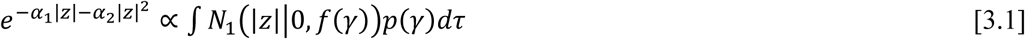

Where the variances are defined as 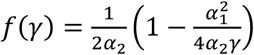. The measurable space in which the variable *z*|*γ* is defined has a unnormalized density function given by the Gaussian pdf *p*(*z*|*γ*) = *N*_1_(|*z*||0, *f*(*γ*)) and its variance *f*(*γ*) is dependent on the random variable *γ* which has Truncated Gamma pdf.

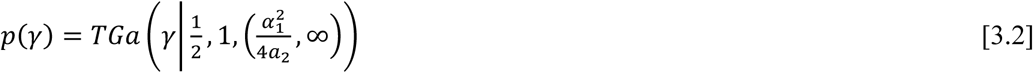

So, the measure in the space product of *z* and *p*(*γ*) is had density represented as an unnormalized product of Gaussian and Gamma densities.

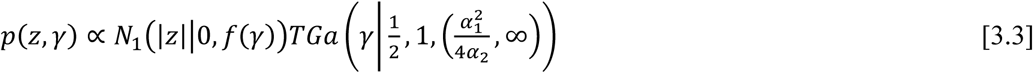

### SI-4 Tensor spatio-frequency prior probabilities of the group Hierarchical Elastic Net

We introduce tensor group penalization on the 3D cartesian space of generators, samples and frequencies with:

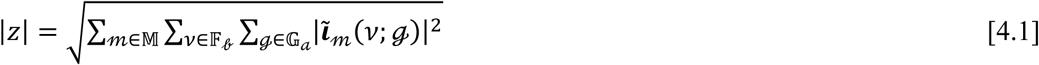

where

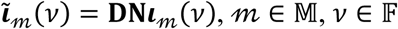

𝒶 refers to a specific Gray matter area 𝒶 ∈ 𝔸

𝒷 refers to a specific frequency band 𝒷 ∈ 𝔹

Then the transformed prior of the parameters is described analytically by the following distribution

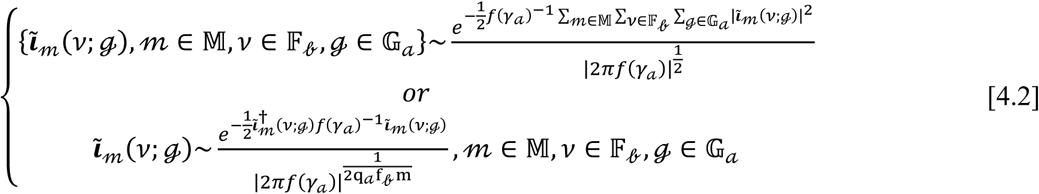

where 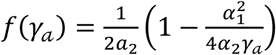 and 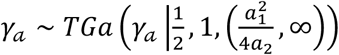

In vector form they are expressed as

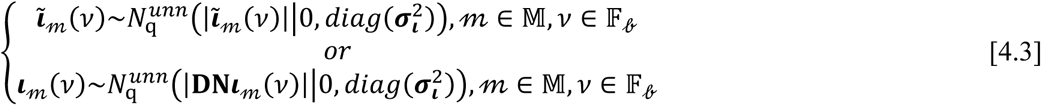

where

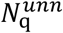 is the unnormalized distribution 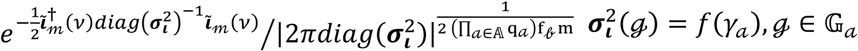, 𝒶 ∈ 𝔸 are the variances

The full vector Bayesian model is as follows:

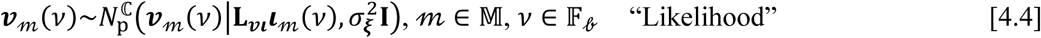

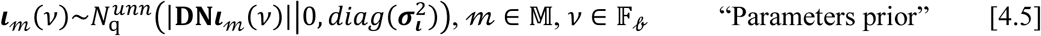

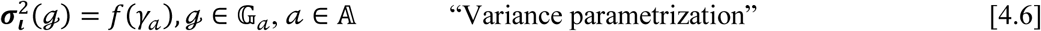

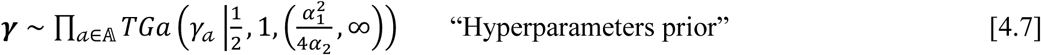

### SI-5 Parameters posterior probability

For the joint distribution of data and parameters *p*(***v***_𝓂_(*v*), ***ι***_𝓂_(*v*)) the following factorization holds, for simplicity we avoid the use of argument for frequency *v* and samples *m*:

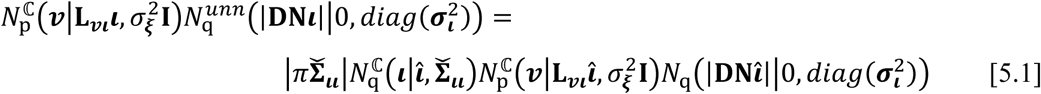

The quantities 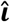 (posterior mean) and 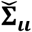 (posterior covariance) are defined as follows

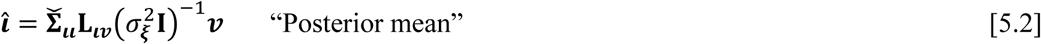

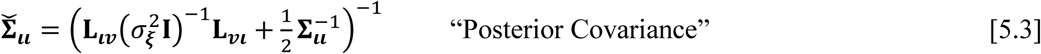

where 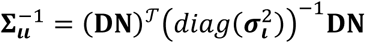

This can be demonstrated by writing their distributions explicitly.

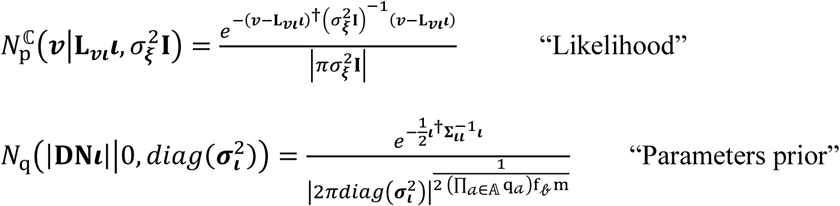

The form of the resultant distribution can be found by analyzing the terms that depend on the parameters (exponential argument) in the formula above

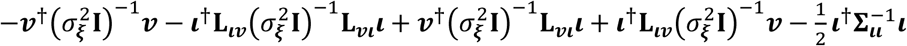

Reorganizing 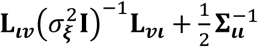 of terms 2 and 5 to render 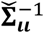 and 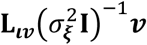 in terms 3 and 4 to render 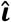

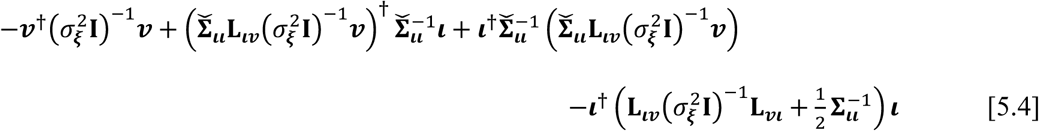

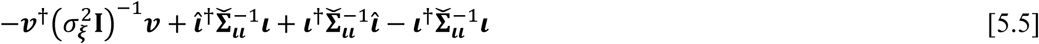

Completing with the term 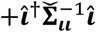

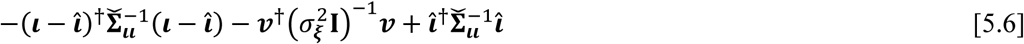

Completing with the terms 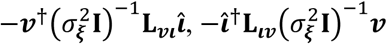 and 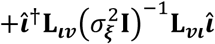 we obtain:

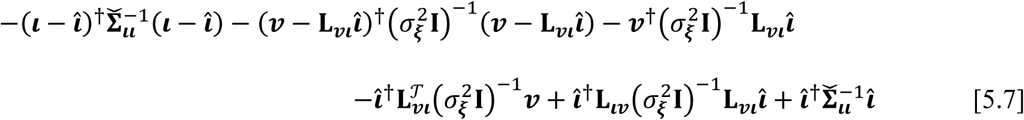

Then terms 3, 4 and 6 can be reorganized into 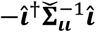 since 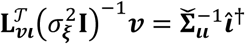 to obtain:

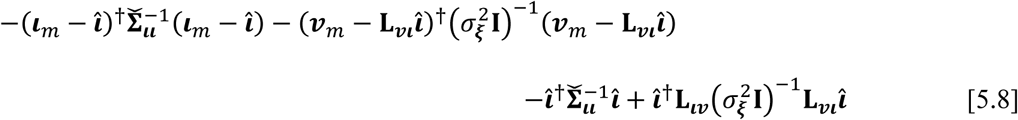

But combining terms 3 and 4 yields 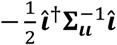

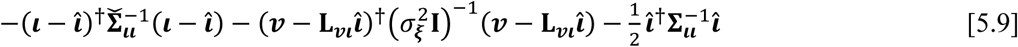

From this, it holds that:

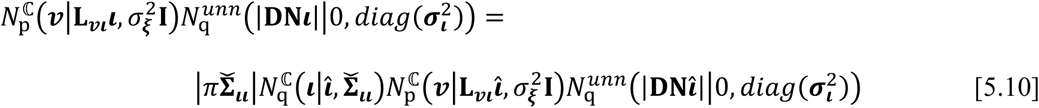

### SI-6 Empirical Bayes and hyperparameters posterior probability

From the joint distribution *p*({(***v***_𝓂_(*v*), ***ι***_𝓂_(*v*)), 𝓂 ∈ 𝕄, *v* ∈ 𝔽_𝒷_}) we can derive iteratively and approximated representation the Type II-Likelihood *p*^(*k*)^({***v***_𝓂_(*v*), 𝓂 ∈ 𝕄, *v* ∈ 𝔽_𝒷_}):

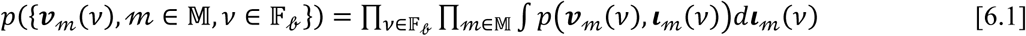

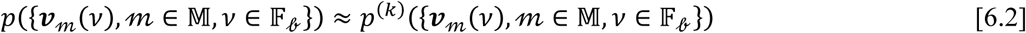

where

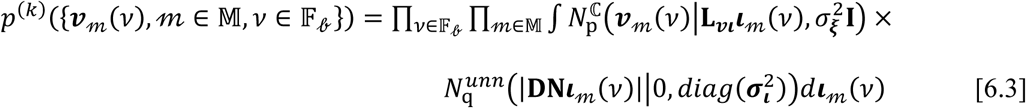

The analysis of section 5 yields

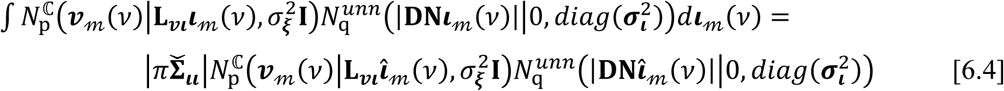

Then iteratively upon fixed values of the parameters 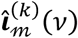 the Type II-Likelihood is expressed as:

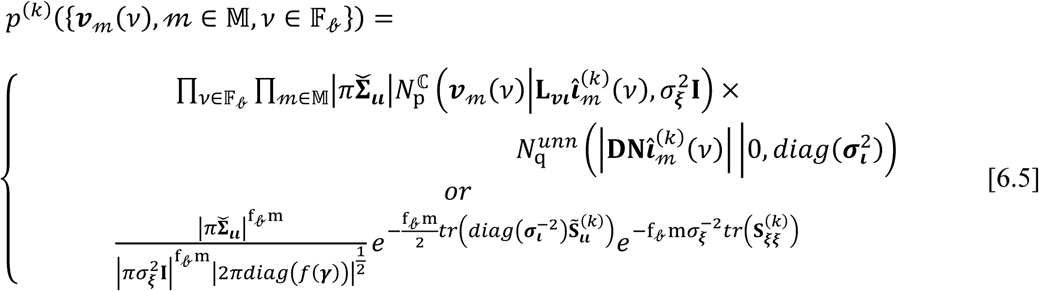

Where the covariances 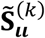and 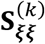 are

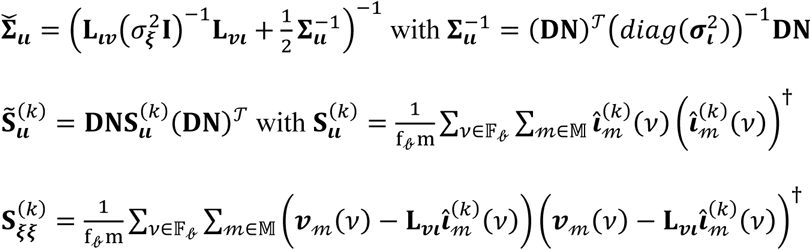

#### Parameter estimators

The parameters 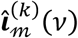 are determined in the previous iteration in terms of the iteratively linear source transfer operator 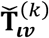

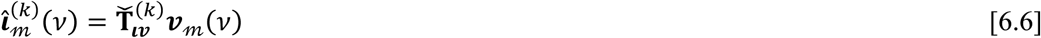

The source transfer operator is defined upon fixed values of the hyperparameters:

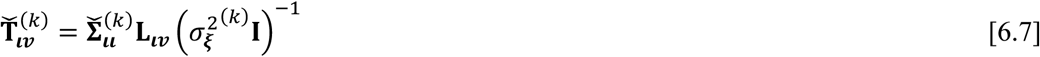

where

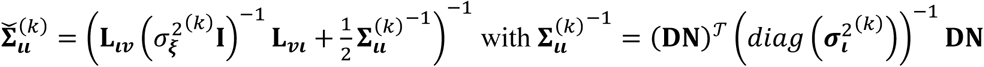

This simplifies the computations of 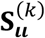 expressed as:

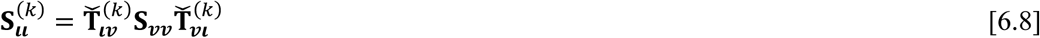

where

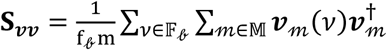

### SI-7 Hyperparameter estimators

The estimation formulas can be derived by applying maximum posterior of the combined Type II Likelihood and priors. To do so we reformulate the targeted hyperparameters as follows:

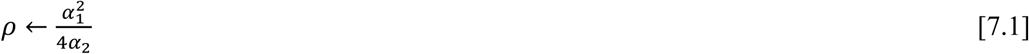

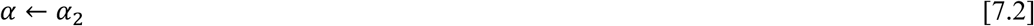

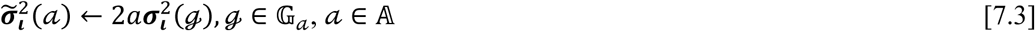

#### Variances

First the estimator of variances 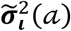 can be computed from the stationary values of the expression:

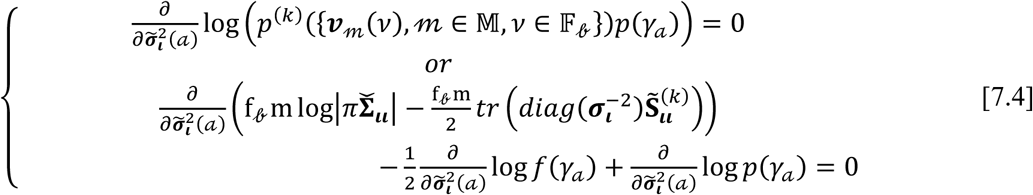

Due to the chain rule of matrix derivatives the first term can be expressed in a close form, see preposition 2.3.2b in [Paz-Linares, 2017].

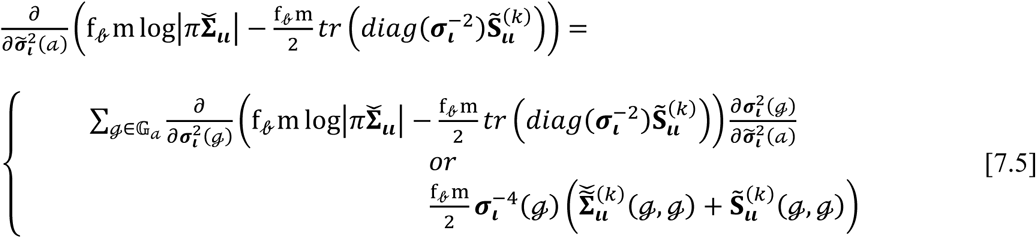

where

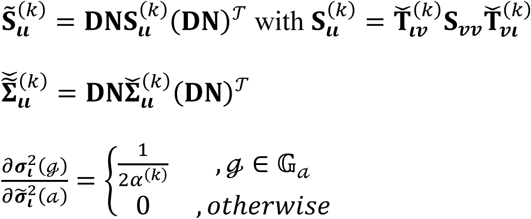

Yielding:

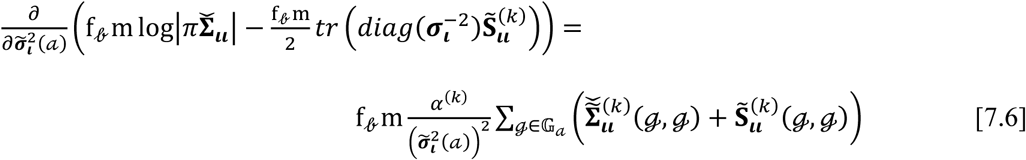

The second and third term derivatives are

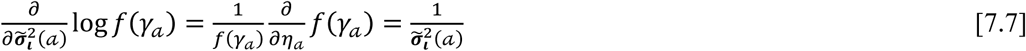

where

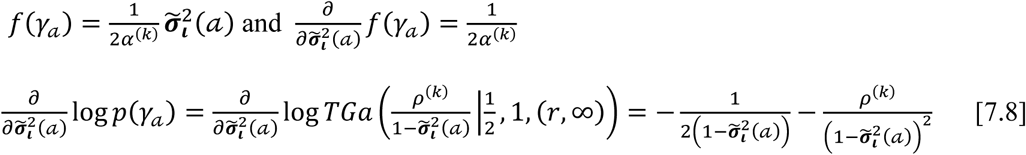

Substituting the derivatives, we obtain:

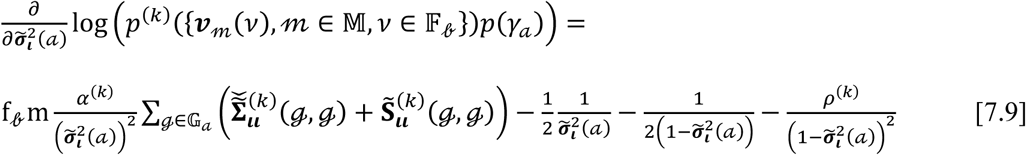

But since it holds that 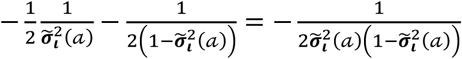 the expression is much more compact:

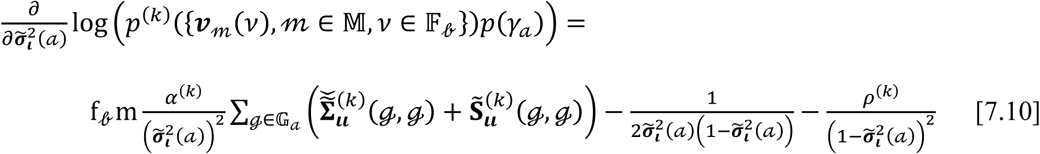

Using the auxiliary quantity *η*_𝒶_ so that 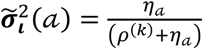 we obtain:

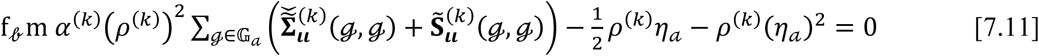

Therefore, the only possible solution for the conditions set by the problem statement and estimator can be obtained from *η*_𝒶_ with:

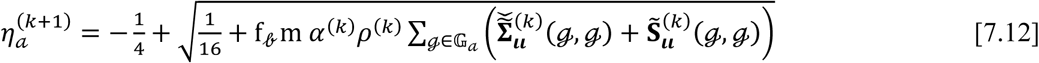

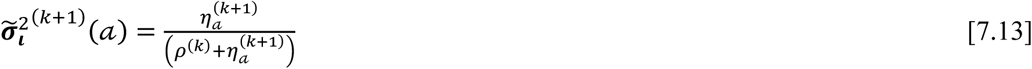

#### Regularization parameters ρ and α

The estimator of the regularization parameters *r* and *a* can be computed from the stationary values of the expression below, following the same steps as for the variances, see preposition 2.3.2c in [Paz-Linares, 2017]:

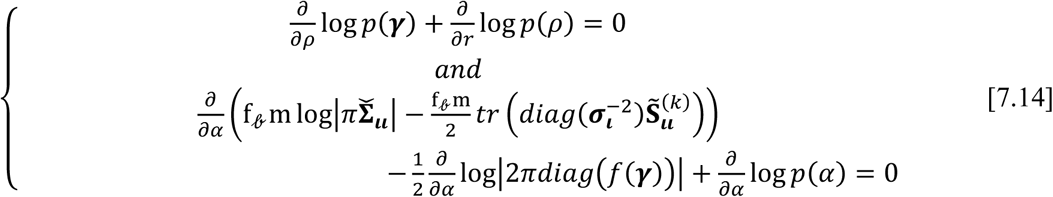

Writing explicitly the *TGa* distribution and using the chain rule in the derivative 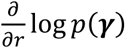:

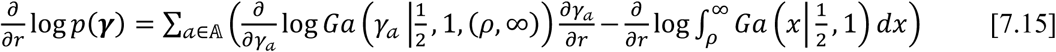

where

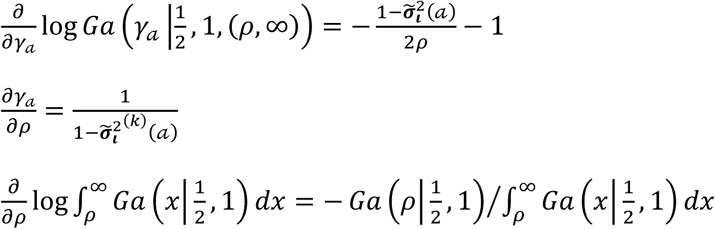

The derivative 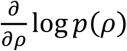 is given by:

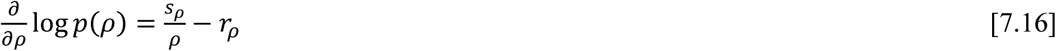

where *p*(*ρ*) ∝ *Ga*(*ρ*|*s*_*ρ*_ + 1, *r*_*ρ*_) with shape (*s*_*ρ*_ + 1) and rate *r*_*ρ*_

From this we obtain the equation for *ρ*:

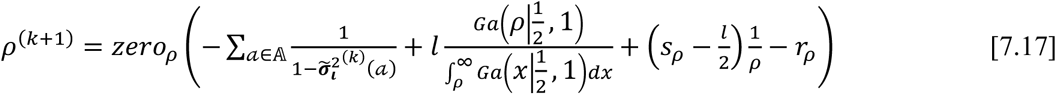

Due to the chain rule of matrix derivatives the first term for the parameter *α* can be expressed as:

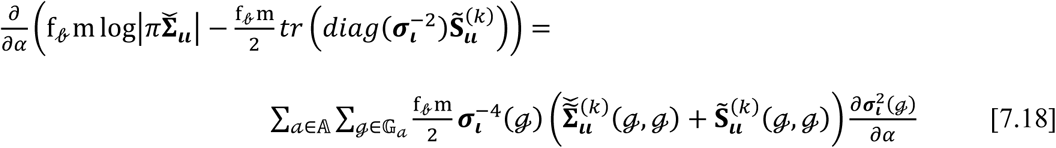

where

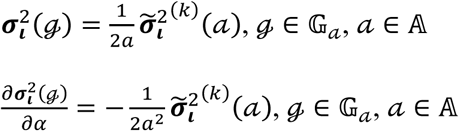

The derivative 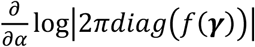 is given by:

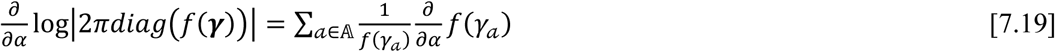

where

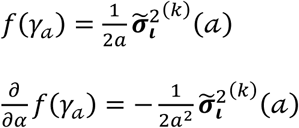

The derivative 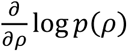 is given by:

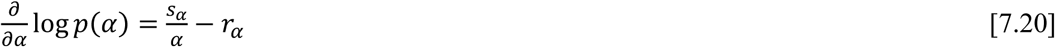

where *p*(*α*) ∝ *Ga*(*α*|(*s*_*α*_ + 1), *r*_*α*_) with shape (*s*_*α*_ + 1) and rate *r*_*α*_

From this we obtain the equation for *α*:

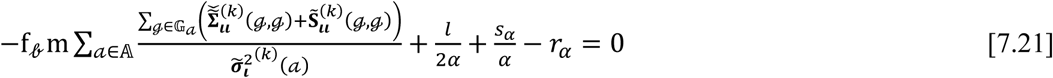

*or*

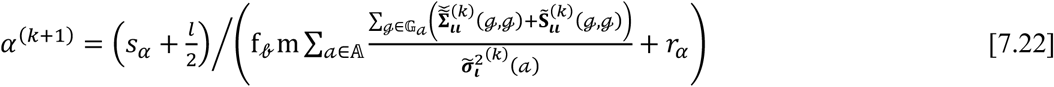

#### Noise parameter 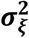

The estimator of the noise parameter 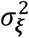 can be computed from the stationary values of the expression below, following the same steps as for the variances, see preposition 2.3.2d in [Paz-Linares, 2017]:

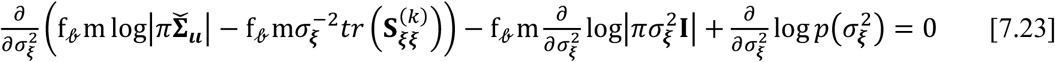

Due to the chain rule of matrix derivatives the first term for the parameter *α* can be expressed as:

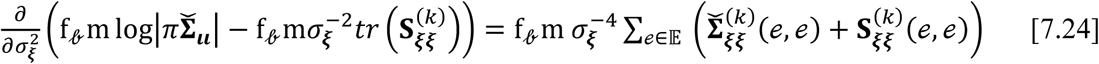

where 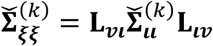 and 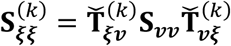 with 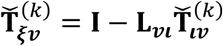

The derivative 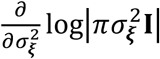 is given by:

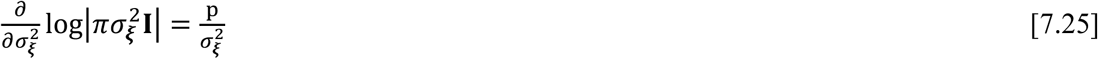

The derivative 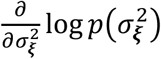 is given by:

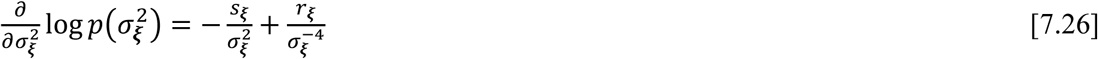

where 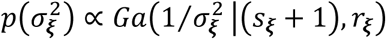 with shape (*s*_***ξ***_ + 1) and rate *r*_***ξ***_

From this we obtain the equation for *α*:

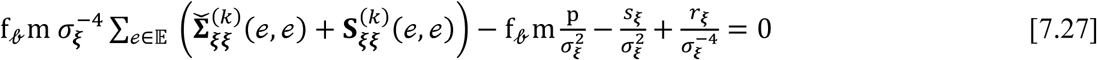

*or*

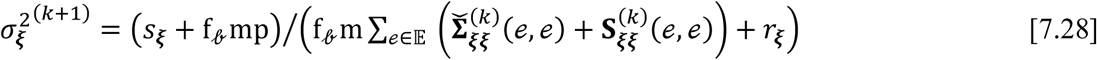

### SI-8 Spectral Structured Sparse Bayesian Learning with multiple priors (sSSBL++)

|  |  |
| --- | --- |
| **Inputs** | $\mathbf{L}_{\iota v}$ (Lead Field), *scaleL* (scale factor for $\mathbf{L}_{\iota v}$), $\{\mathbf{S}_{vv}(\nu), \nu \in \mathbb{F}_\ell\}$ (Observed cross-spectrum), *scaleS* (scale factor for $\mathbf{S}_{vv}$), $p$ (Number of sensors), $f_\ell$ (Band size), $m$ (Sample size), $\mathbf{D}$ (Laplacian) $\mathbf{N}$ (Normal directions), $\{\mathbb{G}_a, a \in \mathbb{A}\}$ (parcels), $l$ (Parcellation size), $s_\xi$ (noise shape), $r_\xi$ (noise ratio), $s_\alpha$ (alpha shape), $r_\alpha$ (alpha ratio), $s_\rho$ (rho shape), $r_\rho$ (rho shape), *maxiter* (Maximum number of iterations) |
| **Outputs** | $\mathbf{T}_{\iota v}$ (Source transfer operator), $\tilde{\mathbf{\Sigma}}_u$ (Source posterior covariance), $\{\mathbf{S}_u(\nu), \nu \in \mathbb{F}_\ell\}$ (Source covariance or cross-spectrum), |
| **Initial values** | $\alpha^{(0)}, \rho^{(0)}, \sigma_\xi^{2(0)}$ and $\sigma_t^{2(0)}$<br>$\mathbf{S}_{vv} = \frac{1}{f_\ell} \sum_{\nu \in \mathbb{F}_\ell} \mathbf{S}_{vv}(\nu)$ |
| | $\mathbf{L}_{uv} = \mathbf{L}_{uv}/scale\mathbf{L}$<br>$\mathbf{S}_{vv} = \mathbf{S}_{vv}/scale\mathbf{S}$ |
| **START** |  |
| **FOR** $k = 1 \dots maxiter$ | |
| **Parameters** |  |
| **step 1** | $\boldsymbol{\Sigma}_u^{(k)-1} = (\mathbf{DN})^T \left( diag \left( \sigma_i^{2(k)} \right) \right)^{-1} \mathbf{DN}$ |
| **step 2** | $\widetilde{\boldsymbol{\Sigma}}_u^{(k)} = \left( \mathbf{L}_{uv} \left( \sigma_\xi^{2(k)} \mathbf{I} \right)^{-1} \mathbf{L}_{vi} + \frac{1}{2} \boldsymbol{\Sigma}_u^{(k)-1} \right)^{-1}$ |
| **step 3** | $\widetilde{\widetilde{\boldsymbol{\Sigma}}}_u^{(k)} = \mathbf{DN} \widetilde{\boldsymbol{\Sigma}}_u^{(k)} (\mathbf{DN})^T$ |
| **step 4** | $\widetilde{\boldsymbol{\Sigma}}_{\xi\xi}^{(k)} = \mathbf{L}_{vi} \widetilde{\boldsymbol{\Sigma}}_u^{(k)} \mathbf{L}_{iv}$ |
| **step 5** | $\widetilde{\mathbf{T}}_{iv}^{(k)} = \widetilde{\boldsymbol{\Sigma}}_u^{(k)} \mathbf{L}_{uv} \left( \sigma_\xi^{2(k)} \mathbf{I} \right)^{-1}$ |
| **step 6** | $\widetilde{\mathbf{T}}_{\xi v}^{(k)} = \mathbf{I} - \mathbf{L}_{vi} \widetilde{\mathbf{T}}_{iv}^{(k)}$ |
| **step 7** | $\mathbf{S}_u^{(k)} = \widetilde{\mathbf{T}}_{iv}^{(k)} \mathbf{S}_{vv} \widetilde{\mathbf{T}}_{vi}^{(k)}$ |
| **step 8** | $\mathbf{S}_{\xi\xi}^{(k)} = \widetilde{\mathbf{T}}_{\xi v}^{(k)} \mathbf{S}_{vv} \widetilde{\mathbf{T}}_{v\xi}^{(k)}$ |
| **Hyperparameters** |  |
| **FOR** $a = 1 \dots l$ | |
| **step 9** | $\eta_a^{(k+1)} = -\frac{1}{4} + \sqrt{\frac{1}{16} + f_\theta m \alpha^{(k)} \rho^{(k)} \sum_{\mathcal{G} \in \mathbb{G}_a} \left( \widetilde{\widetilde{\boldsymbol{\Sigma}}}_u^{(k)}(\mathcal{G}, \mathcal{G}) + \widetilde{\mathbf{S}}_u^{(k)}(\mathcal{G}, \mathcal{G}) \right)}$ |
| **step 10** | $\widetilde{\sigma}_i^{2(k+1)}(a) = \frac{\eta_a^{(k+1)}}{\left( \rho^{(k)} + \eta_a^{(k+1)} \right)}$ |
| **step 11** | $\sigma_i^{2(k+1)}(\mathbb{G}_a) = \frac{1}{2\alpha^{(k)}} \widetilde{\sigma}_i^{2(k)}(a)$ |
| **END** |  |
| **step 12** | $\rho^{(k+1)} = \text{zero}_\rho \left( -\sum_{a=1}^l \frac{1}{1-\hat{\sigma}_t^{2(k)}(a)} + l \frac{Ga(\rho\vert \frac{1}{2}, 1)}{\int_\rho^\infty Ga(x\vert \frac{1}{2}, 1)dx} + (s_\rho - \frac{l}{2})\frac{1}{\rho} - r_\rho \right)$ |
| **step 13** | $\alpha^{(k+1)} = (s_\alpha + \frac{l}{2}) / \left( f_\ell m \sum_{a=1}^l \frac{\sum_{\mathcal{G} \in \mathbb{G}_a} (\tilde{\Sigma}_u^{(k)}(\mathcal{G}, \mathcal{G}) + \mathbf{S}_u^{(k)}(\mathcal{G}, \mathcal{G}))}{\hat{\sigma}_t^{2(k)}(a)} + r_\alpha \right)$ |
| **step 14** | $\sigma_\xi^{2(k+1)} = (s_\xi + f_\ell \text{mp}) / (f_\ell m \sum_{e=1}^p (\tilde{\Sigma}_{\xi\xi}^{(k)}(e, e) + \mathbf{S}_{\xi\xi}^{(k)}(e, e)) + r_\xi)$ |
| **END** |  |
| **Converged estimators** |  |
| **step 15** | $\mathbf{T}_{lv} = \tilde{\mathbf{T}}_{lv}^{(\text{maxiter})}$ |
| **step 16** | $\tilde{\Sigma}_u = \tilde{\Sigma}_u^{(\text{maxiter})}$ |
| **step 17** | $\mathbf{S}_u(v) = \mathbf{T}_{lv} \mathbf{S}_{vv}(v) \mathbf{T}_{lv} \text{ for } v \in \mathbb{F}_\ell$ |
| **END** |  |

#### Algorithm interpretation

The estimation procedure is based on maximum posterior analysis extended from [Paz-Linares et al 2017]. The parameters *ρ* (step 12) and *α* (step 13) tune the global level of sparsity by balancing the contribution of the group Hierarchical Elastic Net of step 9. The selection of specific neural causes is controlled by 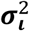 (step 11), which does it for the grouped frequency components *v* ∈ 𝔽_𝒷_, **m** samples and generators into an area **ℊ** ∈ **𝔾**_𝒶_. Thus, allowing to identify the spatial signature of the band robustly and area robustly with the information of all samples.

#### Algorithm statistics

The sSSBL or sSSBL++ allows screening out the neural space by thresholding the posterior distribution statistic: the ratio of the posterior mean and posterior variances. After convergence the estimated source activity can be thresholded by means of an unbiased statistic: this is due to the posterior distribution of source activity 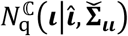 of formula [5.1], where the quantities 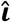 (posterior mean) and 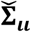 (posterior covariance) are defined as follows:

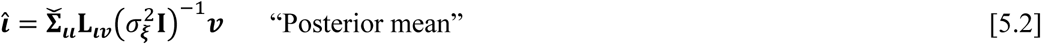

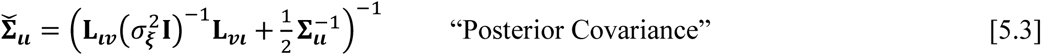

In this distribution, *diag*(**S**_***ιι***_) is the posterior mean and 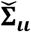 the posterior covariance. The z-statistic for the analysis of variance has the following form: 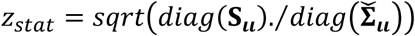. A plausible way to screen out the active sources is to extract the set of nodes 𝒥 that return a value of the z-statistic greater than 1: **𝔾**_0_ = {***ι***: *z*_*stat*_*ι*_ ≥ 1}.

### SI-9 Synthetic EEG

To evaluate the performance of the sSSBL we also use typical synthetic EEG data simulations (***v***_𝓂_, 𝓂 = 1…m) corresponding distributed four sources patches with four different spectral components, see an instance of the “Ground True” in next Figure. Source activity is defined as random complex vectors (***ι***_𝓂_, 𝓂 = 1…m) multivariate Gaussian engine with random Hermitic covariance matrix Σ_***ιι***_. For more realism the covariance is defined from its inverse or precision matrix **Θ**_***ιι***_ a graphical connectivity model. Synthetic data samples (***v***_𝓂_, 𝓂 = 1…m) and its corresponding noisy empirical covariance matrix **S**_***vv***_ were used to evaluate the ESI method performances. To avoid the inverse crime, we used different Lead Field for the inversion to the one used for simulations purpose. The data was corrupted with noise at both levels: (i) sensor (instrumental) noise 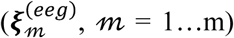 sampled from an univariate Gaussian engine (ii) source (biological) noise 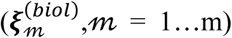 obtained from the Lead Field of an univariate Gaussian engine defined as above. Both types of noise were energy normalized and scaled to a fraction of the EEG signal to achieve the specified signal-to-noise ratio coefficient. See pseudocode for generating synthetic data in next session. For comparison purpose, we use the well-stablished ESI methods MNE and eLORETA.

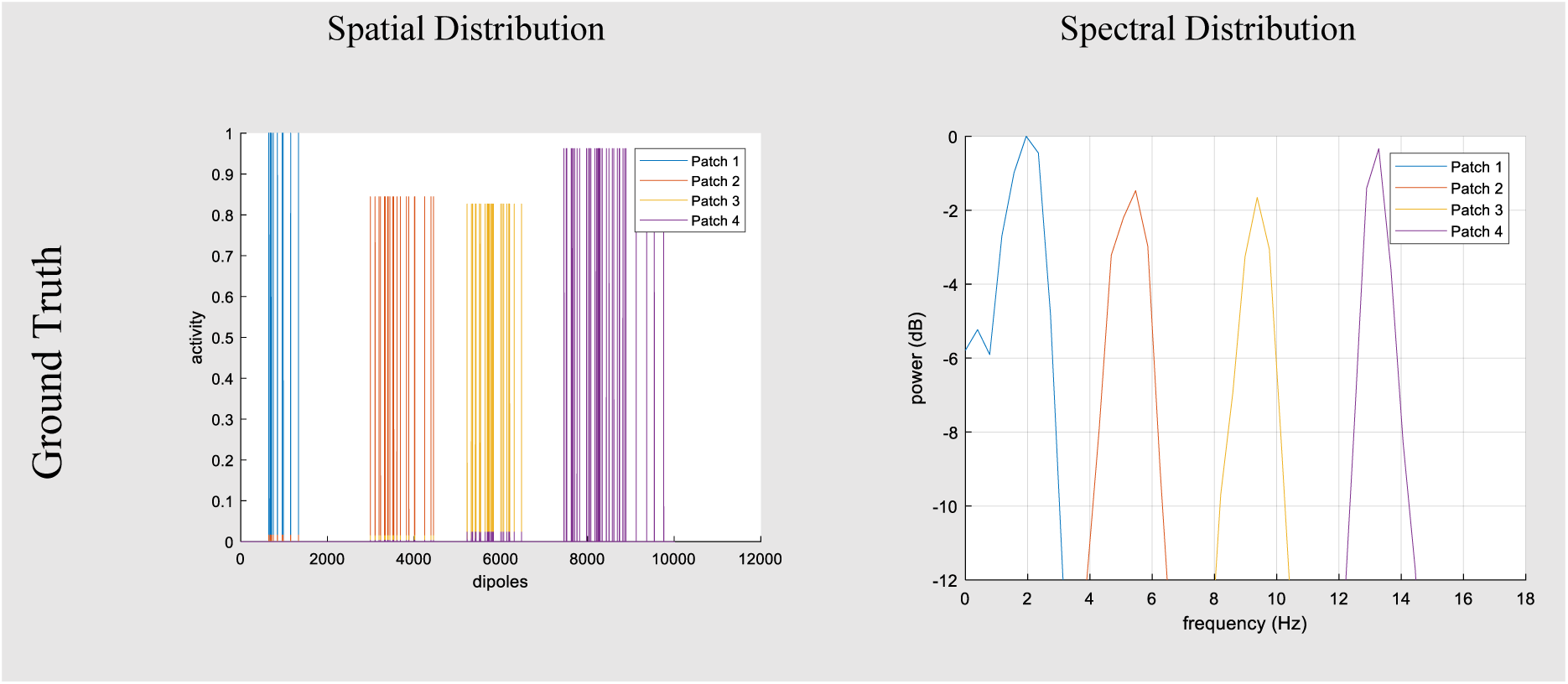

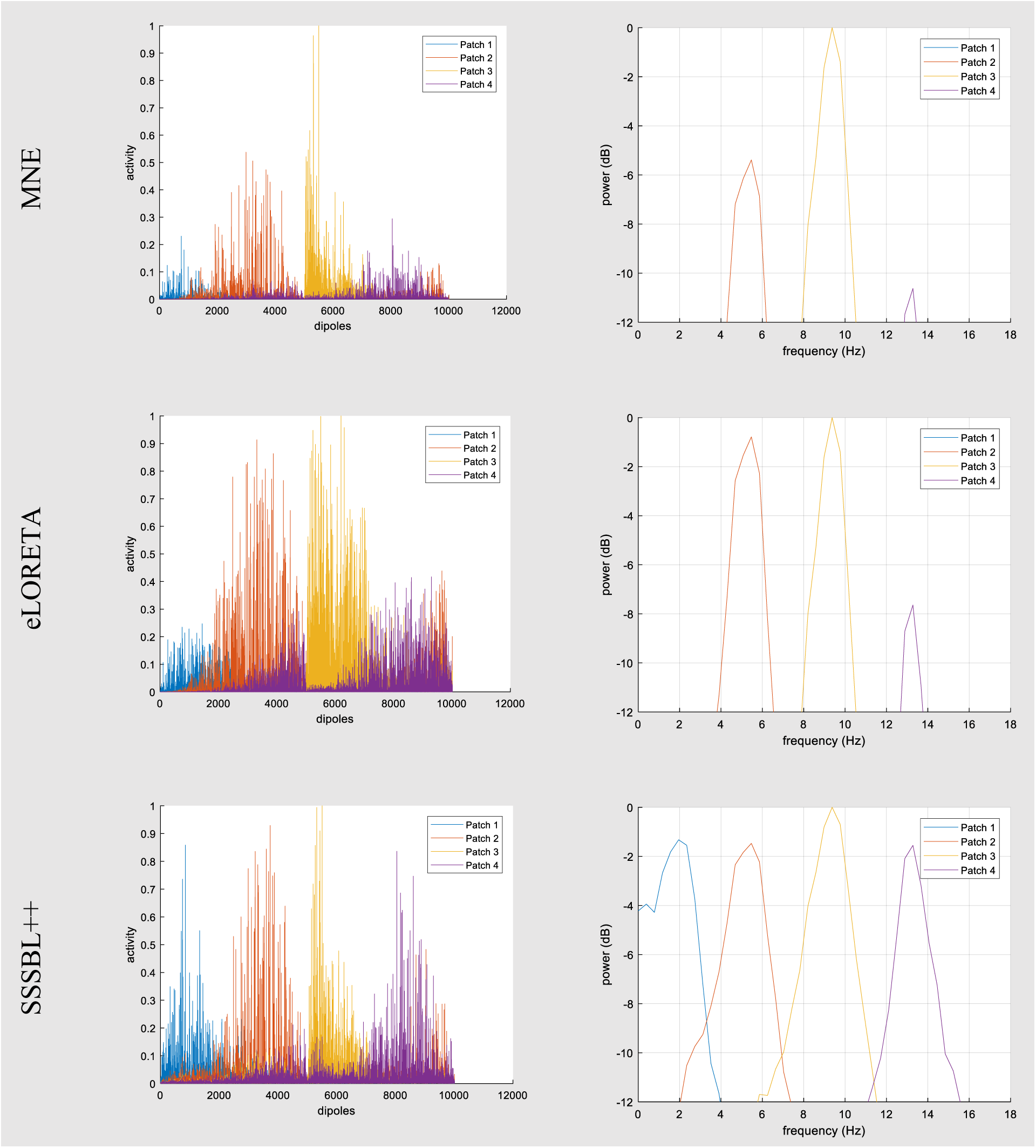

***Results for three ESI methods (MNE, eLORETA and sSSBL++). Simulation framework was based on four patches distributed at cortical surface with different spectral activity.***

For an evaluation of sSSBL++ inference framework, one-hundred random trials of patches centroids were generated. Every random trial included three patches with spatially distributed activity over thirty nodes correspondingly. The synthetic data empirical covariance matrix was obtained from four-hundred samples of simulated source activity and noise. The signal to noise ratio was adjusted to four different levels of noise: none (∞dB), low (19dB), medium (7dB), and high (0dB), as is showed in the next Table.

***EMD MEASURE IN SOURCE LOCALIZATION PERFORMANCE FOR THREE ESI METHODS: MNE, ELORETA, AND SSSBL++. MEAN AND STANDARD DEVIATION ARE SHOWED FOR ONE HUNDRED TRIALS WITH FOUR DIFFERENT LEVELS OF NOISE: NONE (∞DB), LOW (19DB), MEDIUM (7DB) AND HIGH (0DB).***

| NOISE LEVEL \ ESI | MNE | eLORETA | SSSBL++ |
| --- | --- | --- | --- |
| NONE | $8.13 \pm 0.64$ | $12.01 \pm 2.14$ | $5.99 \pm 0.66$ |
| LOW | $14.92 \pm 0.58$ | $20.39 \pm 3.37$ | $11.22 \pm 0.85$ |
| MEDIUM | $14.86 \pm 0.57$ | $41.37 \pm 4.81$ | $10.91 \pm 0.58$ |
| HIGH | $14.89 \pm 0.64$ | $51.65 \pm 2.18$ | $10.92 \pm 0.62$ |

### SI-10 Pseudocode for synthetic data generation

|  |  |
| --- | --- |
| **Inputs** | **L** (Lead Field), **G** (Generator space from a discretization of the Gray Matter), **E** (Sensor space for EEG/MEG), **p** (Number of sensors), **q** (Number of sources), **m** (Sample size), $\sigma_{\xi}$ (Sensors' noise) |
| **Outputs** | **S<sub>u</sub>** (Synthetic source empirical covariance matrix), **S<sub>vv</sub>** (Synthetic noisy data empirical covariance matrix) |
| **START** |  |
| **Activity** |  |
| **step 1** | $\Re(\Theta_u) \leftarrow randn(q, q)$ Real part in subspace **G** |
| **step 2** | $\Re(\Theta_u) \leftarrow (\Re(\Theta_u) + \Re(\Theta_u)^T)/2$ Symmetric property |
| **step 3** | $\Im(\Theta_u) \leftarrow randn(q, q)$ Imaginary part at subspace **G** |
| **step 4** | $\Im(\Theta_u) \leftarrow (\Im(\Theta_u) - \Im(\Theta_u)^T)/2$ Antisymmetric property |
| **step 5** | $\Theta_u \leftarrow complex(\Re(\Theta_u), \Im(\Theta_u))$ Complex compositing |
| **step 6** | Set to zero elements in $\Theta_u$ with unconnected indexes |
| **step 7** | If $\min(svd_s(\Theta_u)) \leq 0$ then correct singular values to be greater than zero |
| **step 8** | $(\iota_m) \sim N_q^C(\iota_m \vert \mathbf{0}, \Theta_u^{-1}); m = 1 \dots \mathbf{m}$ |
| **step 9** | $\mathbf{S}_u \leftarrow \frac{1}{m} \sum_{m=1}^m \mathbf{u}_m \mathbf{u}_m^\dagger$ |
| **Noise** |  |
| **step 10** | $\xi_m^{(inst)} \sim \text{randn}(p, 1); m = 1 \dots \mathbf{m}$ Instrumental noise |
| **step 11** | $\mathbf{S}_{\xi\xi}^{(inst)} \leftarrow \frac{1}{m} \sum_{m=1}^m \xi_m^{(inst)} \xi_m^{(inst)\dagger}$ Instrumental noise empirical covariance matrix |
| **step 12** | $\xi_m^{(biol)} \sim \text{randn}(q, 1); m = 1 \dots \mathbf{m}$ Biological noise at subspace $\mathbb{G}$ |
| **step 13** | $\xi_m^{(biol)} \leftarrow \mathbf{L} \xi_m^{(biol)}; m = 1 \dots \mathbf{m}$ Biological noise at subspace $\mathbb{E}$ |
| **step 14** | $\mathbf{S}_{\xi\xi}^{(biol)} \leftarrow \frac{1}{m} \sum_{m=1}^m \xi_m^{(biol)} \xi_m^{(biol)\dagger}$ Biological noise empirical covariance matrix |
| **Signal to noise ratio** |  |
| **step 15** | $\mathbf{S}_{vv} \leftarrow \mathbf{L} \mathbf{S}_u \mathbf{L}^T$ Synthetic ideal data empirical covariance matrix |
| **step 16** | $\xi_m^{(inst)} \leftarrow \xi_m^{(inst)} \text{tr}(\mathbf{S}_{vv}) / (\sigma_\xi \text{tr}(\mathbf{S}_{\xi\xi}^{(inst)})); m = 1 \dots \mathbf{m}$ Instrumental noise |
| **step 17** | $\xi_m^{(biol)} \leftarrow \xi_m^{(biol)} \text{tr}(\mathbf{S}_{vv}) / (\sigma_\xi \text{tr}(\mathbf{S}_{\xi\xi}^{(biol)})); m = 1 \dots \mathbf{m}$ Biological noise |
| **step 18** | $\xi_m \leftarrow \xi_m^{(inst)} + \xi_m^{(biol)}; m = 1 \dots \mathbf{m}$ Total Noise |
| **Data** |  |
| **step 19** | $\mathbf{v}_m = \mathbf{L} \mathbf{u}_m + \xi_m, m = 1 \dots \mathbf{m}$ Synthetic Noisy Data |
| **step 20** | $\mathbf{S}_{vv} = \frac{1}{m} \sum_{m=1}^m \mathbf{v}_m \mathbf{v}_m^\dagger$ |
| **END** |  |

## Notes

#### Summary of Updates

Some figures and their captions were updated.

https://github.com/egmoreira80/Concurrency_sSSBL-

